# Striping artifact removal in VisiumHD data through nuclear counts modeling

**DOI:** 10.64898/2026.05.04.722591

**Authors:** Paola Malsot, Malte Londschien, Valentina Boeva, Gunnar Rätsch

## Abstract

**Motivation:** 10x Genomics VisiumHD enables spatial transcriptomics at 2 μm × 2 μm resolution but exhibits slide-specific, non-periodic striping artifacts due to lane-width variability. These multiplicative row/column effects distort bin total counts and can bias downstream analyses. The state-of-the-art *destriping* approach is the normalization procedure used as a preprocessing step in `bin2cell`; it applies sequential high-quantile row- then column-wise normalization, which is asymmetric and can introduce edge effects/macro-stripes and distortions of large-scale total-count structure.

**Results:** We propose a statistical destriping approach that leverages nuclei segmentation from the coregistered H&E image. Assuming transcript abundance is constant within each nucleus, we model bin counts with a negative binomial distribution whose mean is a product of a nucleus-specific concentration and row- and column-specific stripe-factors reflecting lane-width variation. We fit all parameters in a generalized linear modeling framework with cross-validated regularization on stripe-factors and iterative dispersion estimation, and use the fitted parameters to correct the observed counts into a destriped image. On synthetic data with known ground truth, our method improves stripe-factor estimation accuracy and reduces error in corrected counts relative to `bin2cell` and `bin2cell`-derived baselines. Across four public VisiumHD slides, it consistently lowers striping intensity while substantially better preserving biological signal present in the large-scale global count structure and avoiding the artifacts introduced by other methods.

**Availability and Implementation:** All source code and links to publicly available data used for this study are available at https://github.com/paolamalsot/destriping-GLM.

**Note:** This manuscript extends the version submitted to Intelligent Systems for Molecular Biology (ISMB) 2026 by describing a new optimization algorithm that yields an approximately tenfold speedup. All plots and benchmarks in this manuscript use the updated implementation.

## 1. Introduction

In 2024, 10x Genomics released a spatial transcriptomics technology called VisiumHD, which considerably increased the resolution of its former analog Visium. The technology ships in the form of a glass slide on which a thin slice of tissue of interest can be deposited and measured. The RNA measurement includes a panel of 18085 genes output on a contiguous grid of 2 μm x 2 μm barcoded squares within a 6.5 mm x 6.5 mm capture area [10x Genomics, 2024b,a, 10x Genomics]. The spatial measurement functions via a two-step hybridization process: first a panel of designed probes (specific to RNA transcripts) hybridizes to biological RNA on the slide, and then this hybridization product hybridizes with spatially barcoded oligos covering the array. As reported in the VisiumHD official product sheet, the width of a row or column in the array varies by up to ±10% [10x Genomics, 2024b]. As a result, the count values in a bin are artificially increased/decreased by this stripe-factor and do not reflect solely RNA transcript concentration in the tissue. This artifact is visible as a non-periodic dense striping pattern in the total counts image [Cervilla et al., 2025] and has been shown to impact downstream analyses such as total counts image segmentation and cell typing [Polahski et al., 2024, Kamel et al., 2025].

Currently, published analyses of VisiumHD data use the `destripe` normalization procedure introduced as a preprocessing step for the `bin2cell` method [Polahski et al., 2024, Kamel et al., 2025, Cerezo-Wallis et al., 2025]. It first corrects the horizontal striping by dividing each bin’s total count by a user-specified quantile (by default 0.99) of the corresponding row. The resulting image is then corrected vertically and finally brought back to original count space by multiplying every bin by the specified quantile of global per-bin count totals. This two-stage procedure leads inherently to an asymmetry in the corrected counts with no biological grounding. Even if the striping effect seems in most places visually removed in the resulting image, we noted that the correction introduced a distortion in the global total-count structure of the original data and some artifacts such as edge effects and macro-stripe effects.

### 1.1. Contributions

In this work, we propose a statistical method to fit and correct the stripe-factors causing the artifact, assuming that the RNA concentration is constant in bins belonging to the same nucleus. We obtain nucleus locations from a nuclei segmentation of the H&E staining image of the same tissue section profiled by VisiumHD; this image is acquired by default and co-registered to the spatial transcriptomics data in standard VisiumHD processing. Our model also takes into account the shot noise and overdispersion typically present in count data by assuming a negative binomial distribution of counts, which is standard in the bioinformatics community [Love et al., 2014]. In the following, we demonstrate that our method effectively destripes the total-counts image while preserving the large-scale structure of the original image. It also alleviates the artifacts introduced by `bin2cell`’s `destripe` correction, such as edge and macro-stripe effects.

## 2. Existing work

Striping artifacts have been observed and more extensively discussed in a variety of imaging fields such as whole-slide microscopy [Pollatou, 2020], remote sensing [Sun et al., 2019], X-ray imaging and computed tomography [Münch et al., 2009]. Existing methods can be roughly distinguished into three categories: frequency-filtering [Chen et al., 2003, Münch et al., 2009], statistical [Gadallah et al., 2000], and variational [Sun et al., 2019]. Methods such as standard-field correction [Seeram, 2019] are to be excluded since we cannot do an empty-tissue measurement with the same slide, and the striping artifact is slide-specific. Among frequency-filtering methods, many assume that the striping is periodic [e.g. Chen et al., 2003], which is not our case, and other methods designed for non-periodic stripes work well if stripes are sparse [Münch et al., 2009]. Variational methods also often assume sparsity in the underlying striping pattern [Sun et al., 2019]. Statistical methods, for instance moment-matching [Gadallah et al., 2000], match the offset and gain of each stripe, and resemble, in essence, the `destripe` normalization proposed in `bin2cell` [Polahski et al., 2024]. They assume that the regions being matched are similar and cause artifacts when this assumption is violated or when the image is not sufficiently large. Wirth et al. [2023] also observed a striping artifact in a microfluidics-based spatial transcriptomics technology called xDBiT and advocates for bin-level normalization, i.e., equalizing counts in every bin. This will, by design, remove all biological signal in the global total counts structure.

## 3. Method

In order to correct the striping artifact present in the raw count data, we first fit *stripe-factors* parameters within a nuclear counts model and then destripe the data with the fitted *stripe-factors*. The outline of the method is laid out in Figure 1.

**Fig. 1.**
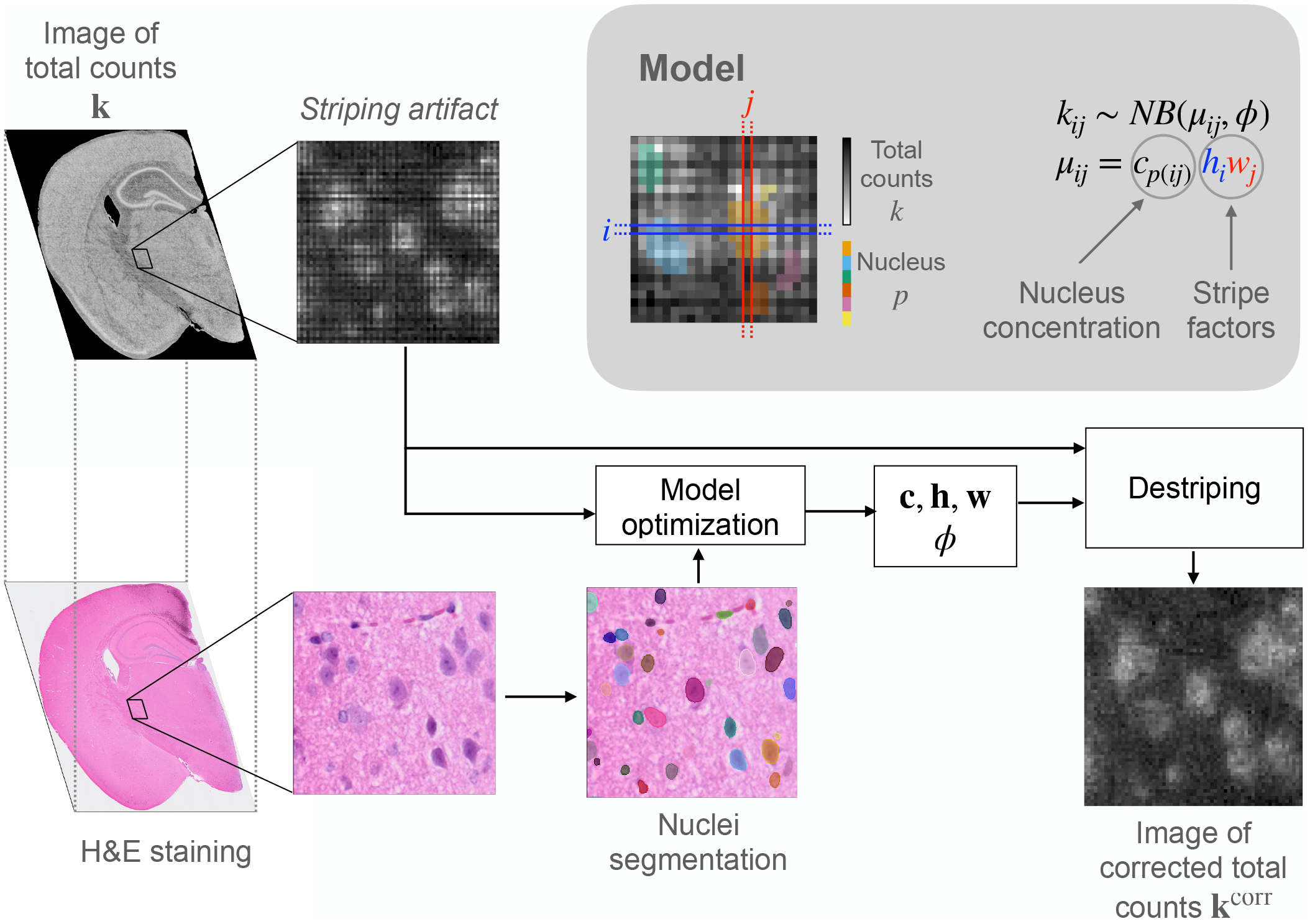
Outline of the proposed method. The raw total-count image **k** exhibits horizontal and vertical striping artifacts. To correct these, our model leverages nuclei segmentation of the H&E staining image. During optimization, we jointly estimate horizontal striping factors **h**, vertical striping factors **w**, nuclei-specific concentration parameters **c**, and a dispersion parameter *ϕ*. The corrected total-count image **k**^corr^ is obtained by destriping the original count image using the fitted model parameters. As illustrated in the upper-right gray panel, the model assumes that the raw count in bin [*i, j* ], *k*_*ij*_, follows a negative binomial distribution with mean μ _*ij*_ = c_*p(I,j)*_ *h*_*i*_ *W*_*j*_ and overdispersion *ϕ*., where *h*_*i*_ and w_j_ denote the striping factors for row *i* and column j, respectively, and *C*_*p*_ represents the concentration of counts within nucleus *p*, with p(*i, j*) indicating the nucleus associated with bin [*i, j*].

### 3.1. Nuclear counts model

According to the manufacturer documentation, the physical width of a lane in the array varies, causing the striping artifact [10x Genomics, 2024b]. Therefore the true size of the bin in row *i* and column *j* equals *h*_*i*_*w*_*j*_, where *h*_*i*_ and *w*_*j*_ are the corresponding horizontal and vertical lane widths respectively. We further assume that the total transcript concentration is homogeneous within nuclei, such that the expected value of total counts in bin [*i,j*], μ_*ij*_ is

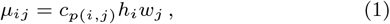

where *c*_*p*_ is the total transcript concentration in nucleus *p* and *p(i,j)* denotes the nucleus to which bin [*i,j*] belongs. As is standard in the bioinformatics community [Love et al., 2014], we assume a negative binomial distribution for count data with dispersion parameter *ø*, such that the following mean-variance relationship holds:

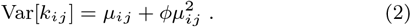

The negative binomial model can be rewritten as a generalized linear model (GLM) with a log-link between the mean of the distribution *μ*_*ij*_ and the linear predictor (*Xβ*)_*ij*_:

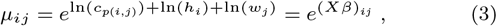

with *β* being a concatenation of the parameters 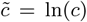, 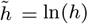 and 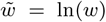 and *X* a carefully chosen design matrix that translates the bins belonging to a particular row, column and nucleus index. For ease of exposition, in slight abuse of notation, we index (Xβ)_*ij*_ = *log(μ*_*ij*_) with *ij*, even though *Xβ* is a row vector. The negative binomial’s negative log-likelihood, up to factors constant w.r.t. *μ*_*ij*_ is

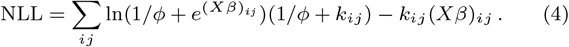

This is convex in *β* and can be minimized with standard solvers. Our model is fitted by minimizing the objective function

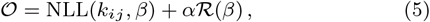

where ℛ (*β*) is a regularization term which can help convergence and generalization and *α ∈* ℝ_+_ is the regularization strength.

Since there is an indeterminacy of scale between the parameters **c, h** and **w**, we alleviate it by imposing mean(**h**) = 1 and mean(**w**) = 1. With this convention the stripe-factors *h*_*i*_, *w*_*j*_ can be interpreted as the actual lane width divided by the typical lane width, and *c*_*p*_ as the counts per typical bin area in the nucleus *p*.

### 3.2. Implementation

#### Nuclei segmentation mask

Nuclei locations are obtained by segmenting the H&E staining image of the tissue slice with `Stardist` [Schmidt et al., 2018] or using an already available segmentation.

#### Fitting the model

We minimize the objective (eq. 5) by block coordinate descent, alternating L-BFGS steps [Liu and Nocedal, 1989] in 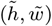 with Newton steps in 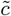. We use the Python library `glum`, which is optimized for fitting generalized linear models with sparse design matrices [Schmidt et al., 2025]. We initialized the *β* parameter in correspondence with *h*_*i*_ = 1, *W*_*j*_ = 1 and 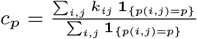.

### Regularization hyperparameter selection

We chose the best regularization parameter *α*with internal crossvalidation among a grid of 10 log-spaced α values between 10^−5^ and 10. Moreover, we compute the solutions for a decreasing sequence of a using warm starts [Friedman et al., 2010]. Finally, the best model is fitted on the whole dataset.

### Dispersion parameter estimation

In order to estimate the dispersion parameter *ϕ*, we alternately:

1. fit the GLM coefficients for a given *ϕ*
2. update *ϕ* with the estimated *μ*_*ij*_ using a similar method as MASS library’s theta.md function [Venables and Ripley, 2002]. In short, it finds *ϕ* by equating the deviance to the residuals’ degrees of freedom until *ϕ* converges.

### 3.3 Destriping the data with the fitted stripe-factors

In order to remove the striping artifact from the data, we perform quantile-mapping from the distribution of striped data *NB (ϕ* _*i,j*_, *ϕ* to the distribution of unstriped data NB*(c*_*p*_(_*i,j*_*), ϕ)* [Robinson and Smyth, 2008]. More specifically, the corrected counts are

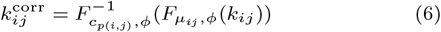

with *F*_*μ,ϕ*_ the cumulative distribution of a negative binomial distribution with parameters *μ* and *ϕ*, and *F*^-**1**^ denoting its inverse, the percentile point function. In the cytoplasm, where c is not estimated, we simply divide the raw counts by the stripe-factors: 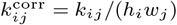.

## 4. Synthetic data generation

As ground truth is unavailable for real data, we construct a synthetic dataset that enables quantitative evaluation of destriping methods.

To reproduce a realistic large-scale total-count structure, we fix the nucleus-to-bin assignment *p(i, j)* to that obtained by segmenting the H&E image of a real VisiumHD mouse brain slide (i.e., the nuclear mask and nucleus identities), and we use nucleus concentrations estimated from the same slide. Synthetic stripe-factors **h** and **w** are generated by sampling row-wise and columnwise effects from a Weibull distribution with parameters fitted to match the empirical distribution of stripe-factors estimated by b2c-sym on the mouse brain slide. We then normalize them such that Σ_*i*_ *h*_*i*_ = *n*_rows_ and *Σ*_*j*_ *W*_*j*_ = *n*_cols_.

We generate gene-specific nuclear counts using a negative binomial model

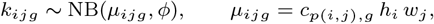

where *C*_*p(i,j),g*_ denotes the mean expression of gene *g* in the nucleus assigned to bin *(i,j)* (estimated from the real mouse brain slide). Non-nuclear bins are set to zero. The dispersion parameter *ϕ*is estimated from the mouse brain slide. Note that the resulting total counts (summed over genes) are approximately negativebinomial. An example of the resulting striped total-count image (zoomed-in region) is shown in Fig. 2A.

**Fig. 2.**
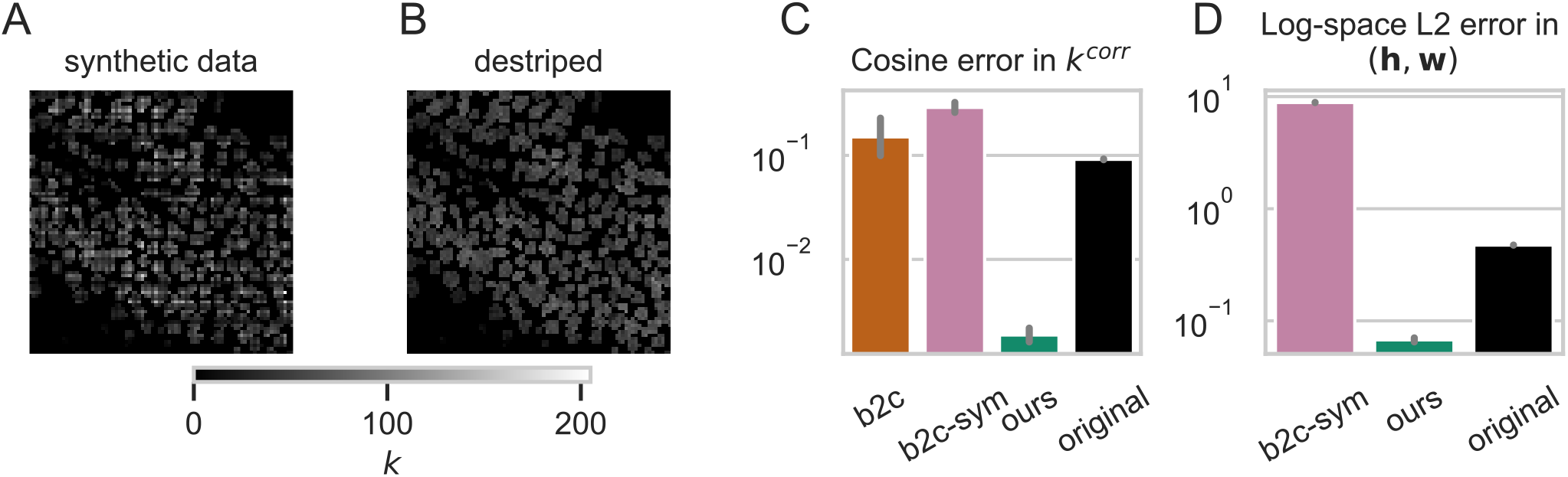
Validation of synthetic data. (A) Illustration of the striping artifact in a region of the synthetic data total-counts image. (B) The same region after correction using ground-truth stripe-factors, where the striping artifact is fully removed. The corrected image containing this region is used as ground-truth reference for panel (C). (C) Comparison of stripe-correction methods using the cosine error of corrected counts, defined as the cosine distance between the corrected counts and the ground-truth reference shown in (B). (D) Comparison of stripe-factor estimation accuracy, measured by the 𝓁_2_ error between the estimated stripe-factors (**h, w**) and the ground truth in log-space. Method b2c is omitted in (D) because it does not explicitly model stripe-factors. Overall, our model achieves superior performance in both stripe-factor estimation (D) and count correction (C). Error bars indicate the standard deviation across three data-generation seeds.

## 5 Evaluation

### 5.1. Baselines

In order to evaluate our destriping method, we compare it to the following baselines.

#### b2c

`bin2cell` sequentially normalizes by row- and column-wise 99^th^ quantiles and rescales by the global 99^th^ quantile [Polafiski et al., 2024]:

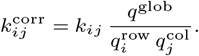

Here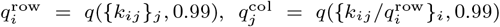,and q^g1ob^ = *q({k*_*ij*_ }_*i,j*_, 0.99).

#### b2c-sym

A symmetric variant that estimates row and column factors directly from the raw image via 99^th^ quantiles and removes them jointly:

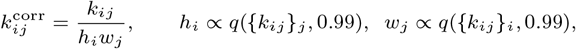

with h and w normalized to have mean 1.

#### bin-level norm

As a reference upper bound on stripe removal, we set all nonzero bins to the median count:

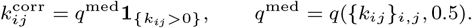

#### Additional baselines

For the interested reader, we include in section S2 of the Supplementary Material several variants of b2c-sym that replace the 99^th^ quantile with the median and/or restrict the estimation to nuclei counts only. We also derive a simple estimator relying on our homogeneity hypothesis for nuclear counts.

### 5.2. Metrics

#### 5.2.1. Synthetic data

On synthetic data, the stripe-factors used for data generation are known and therefore serve as ground truth. Ground-truth corrected counts, denoted *k*^corr,GT^, are obtained by destriping the original count data with ground-truth stripe-factors and nuclei concentrations through quantile mapping (Eq. 6).

The accuracy of the corrected total-count image *k*^corr^ is quantified by the **cosine error** and the **normalized 𝓁**_2_ **error**, defined respectively as the cosine distance and the 𝓁_2_ distance (normalized by the square root of the number of bins) to *k*^corr,GT^.

The accuracy of the estimated stripe-factors **h** and **w** is assessed by comparing them to the ground-truth stripe-factors in log-space. Specifically, we apply the transformation log(x + ∈) with ∈ = 10^-20^ and compute the 𝓁_2_ distance between the transformed estimated and ground-truth stripe-factors, normalized by the square root of the number of lanes. Horizontal and vertical stripe-factor errors are then combined as 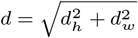.

#### 5.2.2. Real data

On real data, the ground truth is unknown. We therefore define metrics that capture desirable properties of a destriping method.

##### Global structure alteration

Empirically, b2c-derived baselines tend to remove large-scale biological gradients present in real data and create artifacts. To quantify preservation of global structure, we consider the *totalcount profile* along rows and columns, obtained by summing counts per lane and smoothing the resulting curve using a moving average with window size *k* = 100. At this scale, we assume the profile reflects underlying biological structure rather than striping artifacts.

For rows and columns separately, we compute the cosine distance between the smoothed total-count profile of the destriped image and that of the original image. The cosine distance is used to make the metric independent of overall intensity scaling, as we do not wish to penalize methods solely for differences in absolute count magnitude (as observed, for example, with b2c). This choice is further motivated by downstream analyses, which typically rely on relative rather than absolute count differences within a slide. For completeness, we also report in the Supplementary Material a version of the global structure alteration metric based on a normalized Euclidean distance rather than the cosine distance^1^.

##### Striping intensity

To directly quantify the strength of striping artifacts, we introduce a simple striping intensity metric. For horizontal striping, the metric captures systematic differences between vertically adjacent rows by summing bin-wise differences across columns, such that opposing differences cancel out. These row-wise differences are then aggregated into a single score.

Formally, the horizontal striping intensity is defined as

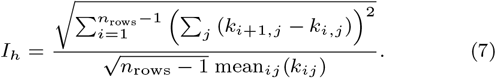

The normalization by 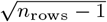 ensures slide-size invariance, while normalization by the mean count makes the metric independent of overall brightness. The inner sum corresponds to differences in total intensity between adjacent rows.

An analogous definition yields the vertical striping intensity *I*_*w*_. The total striping intensity is then given by 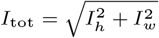.

##### Cytoplasmic striping intensity

In the main text, we report a variant of the striping intensity metric in which the difference *k*_*i+1,j*_ *− k*_*i,j*_ is computed only for pairs of vertically adjacent bins that both belong to the cytoplasm, i.e., for which neither bin is assigned to a nucleus. Restricting the metric to cytoplasmic regions makes the evaluation more stringent for our method, as it excludes the nuclear bins that explicitly inform stripe estimation through the homogeneity assumption. Consequently, strong performance under this metric provides a conservative assessment of destriping quality.

## 6. Results

### 6.1 Results on synthetic data

Figure 2 compares the performance of all methods on simulated data across three data-generation seeds. Our method consistently outperforms the b2c and b2c-sym baselines, yielding lower 𝓁2 error in the log-stripe-factors as well as lower cosine and normalized 𝓁 _2_ errors in the corrected counts *k*^corr^ (see Fig. S4). Given that the simulation model closely matches the assumptions underlying our method, this result is not unexpected. Rather than serving as a definitive benchmark, the simulation primarily validates the correctness and robustness of our optimization procedure. In particular, this experiment allowed us to verify that careful initialization is critical for successful convergence, and that selectively applying an 𝓁_*2*_ regularization to the stripe-related parameters is essential. We use a quadratic regularization term of the form

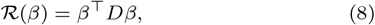

where *D* is a diagonal matrix with nonzero entries corresponding only to the components of *β* associated with log(**h**) and log(**w**), while the nucleus-specific parameters log(**c**) are left unregularized. An ablation study illustrating the impact of this choice is provided in Supplementary Fig. S1. The same supplementary figure also shows that variants of b2c-sym that restrict stripe estimation to nuclear bins and/or replace the 99^th^ quantile with the median improve upon the standard b2c-sym baseline. However, these modifications do not alter the overall ranking, as our method remains consistently superior across metrics. Finally, the simulation benchmark reproduces a known failure mode of b2c-derived baselines: excessive alteration of global structure in the corrected counts. Specifically, b2c and b2c-sym exhibit substantially higher global-structure alteration scores than our method (Supplementary Fig. S1). Moreover, as shown in Supplementary Section S5.1, our correction yields higher celltyping accuracy than the competing baselines.

### 6.2 Results on real data

We evaluated all methods on four publicly available VisiumHD datasets: mouse brain, mouse embryo, human lymph node, and zebrafish head, each consisting of a single slide. Figure 3 summarizes the performance of the destriping methods across these datasets using the two metrics defined above: striping intensity and global structure alteration. Across all datasets, our method consistently achieves substantially lower global structure alteration than the competing approaches, indicating improved preservation of the biological signal contained in the totalcount structure. This conclusion remains unchanged when global structure alteration is quantified using the normalized Euclidean distance instead (Supplementary Fig. S11, S20, S25, and S30). At the same time, it achieves equal or superior reduction in striping intensity on all datasets except the zebrafish head, where the difference relative to b2c is small and comparable. On that dataset, both b2c and b2c-sym introduce large regions of artificially elevated counts, most notably in the lower third and the upper fifth of the slide (see Figure 4). In contrast, our method preserves the original large-scale intensity distribution while effectively reducing the striping visible in the raw data. This is also reflected in a downstream differential-expression analysis on the zebrafish head, where b2c-derived baselines can invert log fold-change directions for a substantial subset of genes, unlike our method (Supplementary Section S5.2). On the human lymph node dataset, our method visibly and quantitatively suppresses the striping artifact more effectively than b2c and b2c-sym, while avoiding the introduction of artifacts such as noisy stripe patterns at the edge of the tissue and large regions of elevated counts (see Figure 5). Detailed results for the mouse brain and mouse embryo datasets are provided in the Supplementary Material. For the mouse brain dataset, b2c and b2c-sym introduce visible distortions, including a horizontal macro-stripe of elevated intensity across the center of the tissue (Supplementary Fig. S6) as well as noisy high-intensity stripes near the bottom edge of the slide (Supplementary Fig. S9). Similarly, on the mouse embryo dataset, b2c and b2c-sym substantially alter the global count structure (Supplementary Fig. S15 and S16), whereas our method better preserves the large-scale count distribution. In addition, a localized region highlights that our method more effectively removes striping than b2c and b2c-sym (Supplementary Fig. S17 and S18). Since our method relies on nuclei segmentation, we also evaluated its robustness to segmentation errors, including missing cells, oversegmentation, and undersegmentation (Supplementary Section S4). These experiments showed that the method is highly robust to missing cells and oversegmentation, and robust to moderate undersegmentation. Only under extreme undersegmentation did performance degrade appreciably, mainly through increased distortion of the global count structure despite effective removal of the striping artifact.

**Fig. 3.**
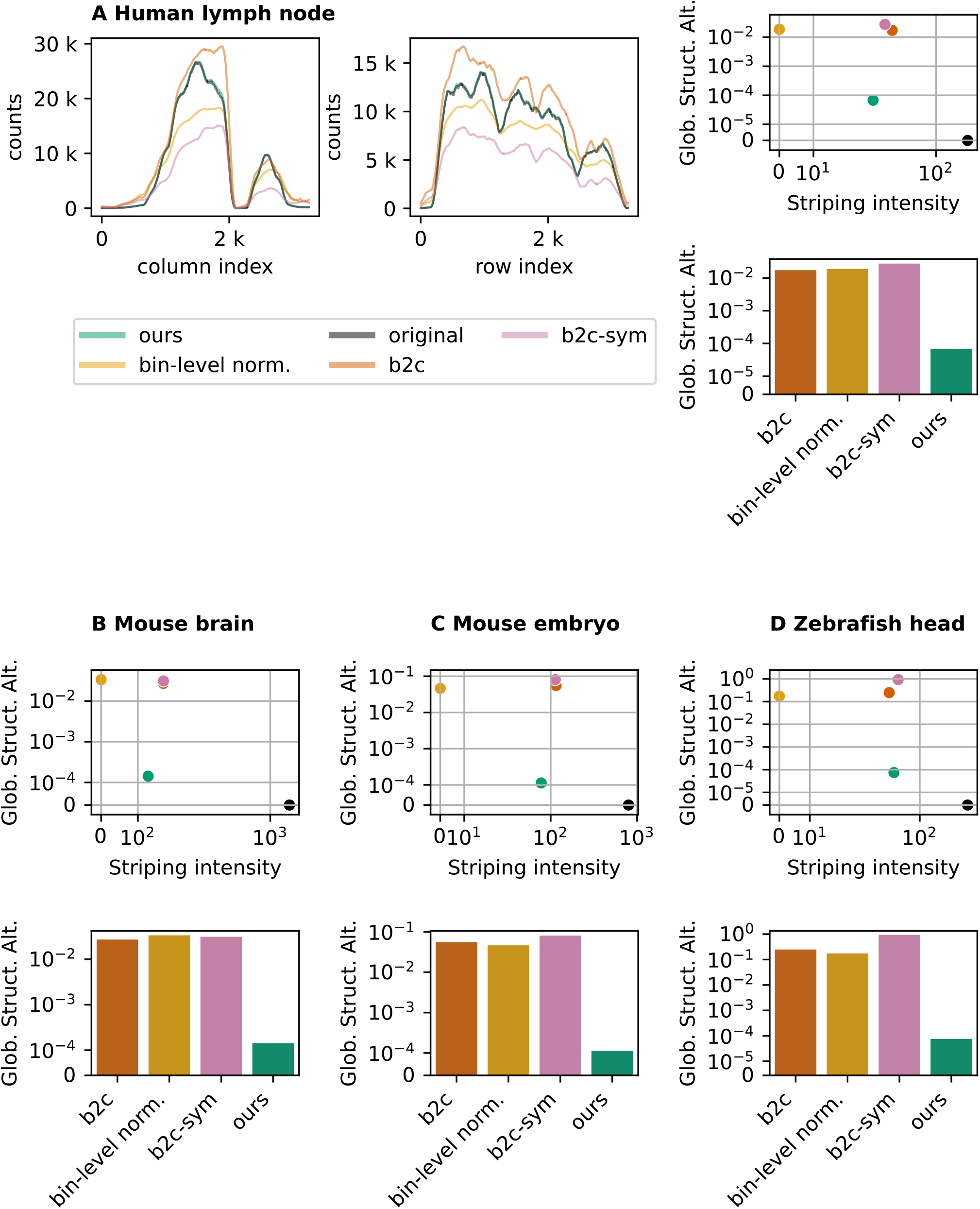
Evaluation of destriping methods on the four experimental datasets: (A) human lymph node, (B) mouse brain, (C) mouse embryo, and (D) zebrafish head. In panel (A, left), we display the smoothed count profiles used to compute the global structure alteration metric (Glob. Struct. Alt.). As desired, our method follows the original profile much closer than the other methods, which is reflected in a lower global structure alteration score (see A-right, bottom). In subpanel (A, upper-right), we show the combination of the striping-intensity and global structure alteration score. Our method achieves as desired lower striping intensity while maintaining the global structure. As a reference, we include bin-level norm which achieves 0 striping intensity by definition, but which considerably alters the global count structure. The superiority of our method is further confirmed in the other datasets (panels B, C and D). Note that in (D), the b2c striping intensity is slightly lower than ours but is counterbalanced with a much higher global count structure alteration.

**Fig. 4.**
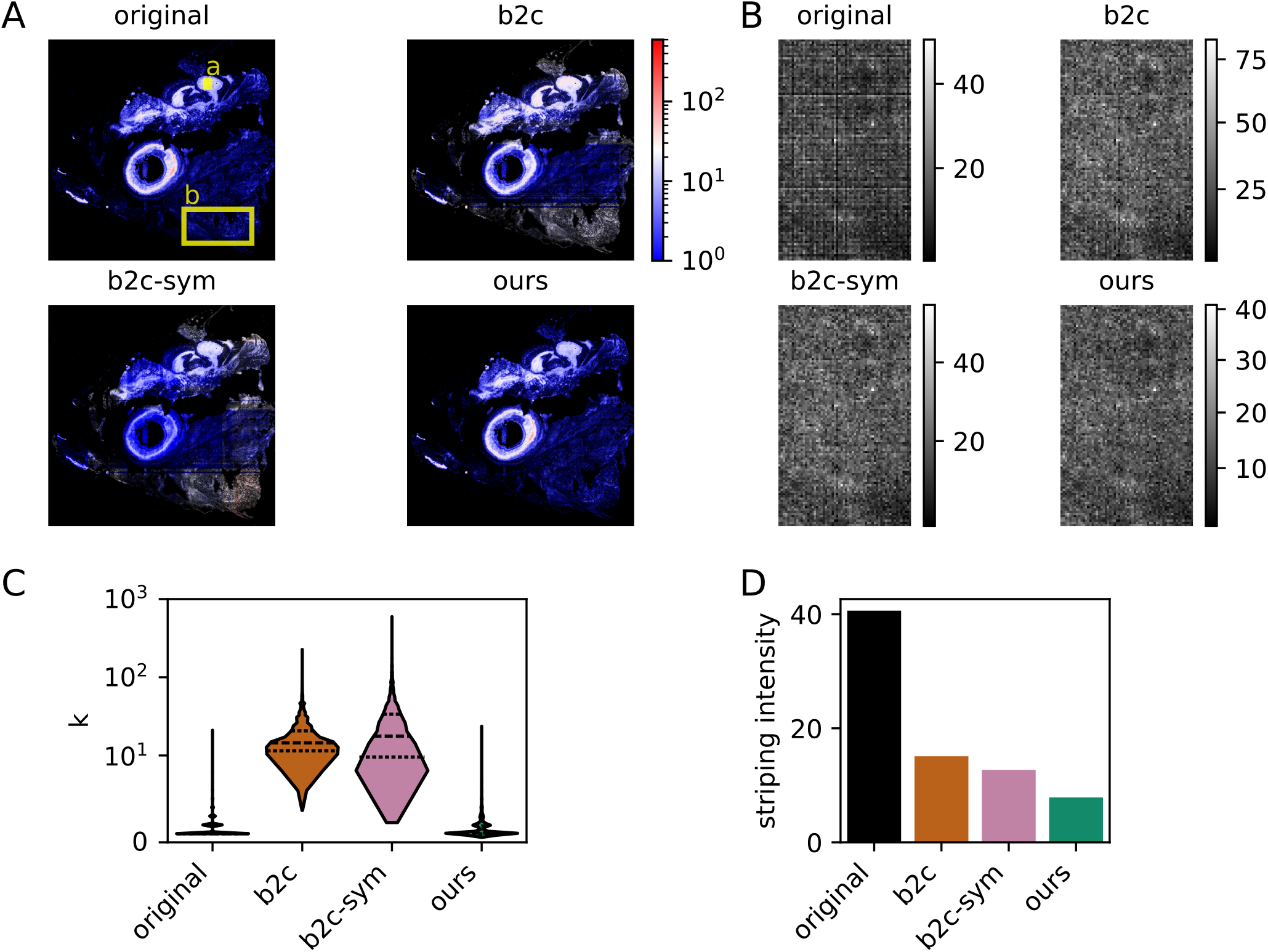
Comparison of destriping methods on the zebrafish head dataset. (A) Entire image of total counts along with its corrections for the methods b2c, b2c-sym and ours. Both b2c and b2c-sym display a three-band structure, with the lower and the upper band having much higher count-values. (C) Count distribution in region b, which is located in the lower band. As expected our count values lie on a comparative scale with original, whereas b2c and b2c-sym show exaggerated values. (B) Zoom of region a for all destriping methods. The destriping looks satisfactory and comparable, if not slightly more pronounced in our method. (D) Comparison of striping intensity on subregion a: indeed, our method achieves lower striping intensity.

**Fig. 5:**
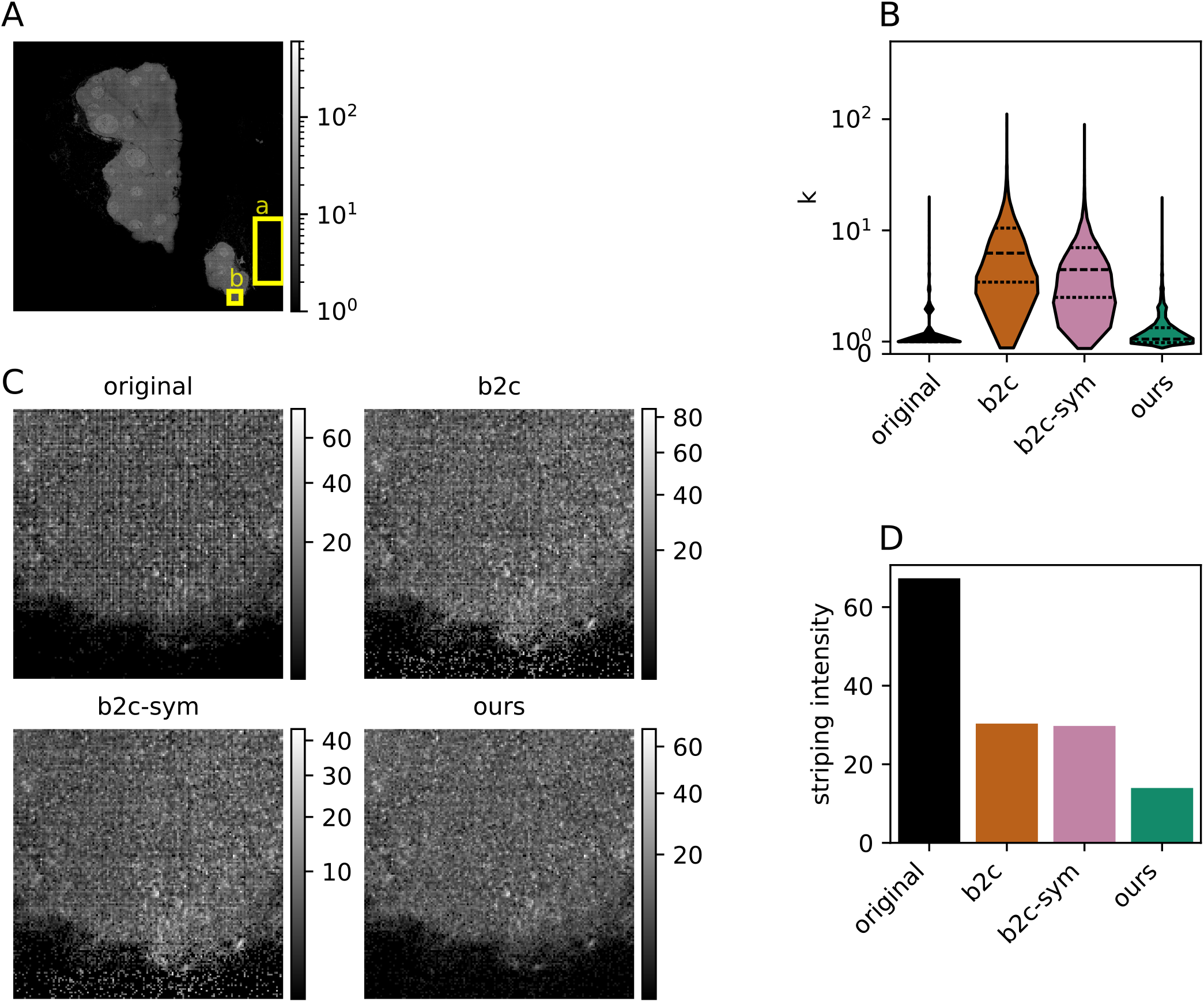
Comparison of destriping methods on the human lymph node dataset. (A) Whole original total counts image, with subregions a and b indicated. (B) Count distribution in region a. b2c and b2c-sym corrections create artificially high values. (C) Zoom on subregion b. b2c and b2c-sym have introduced high-count values in the bottom region, which is dark in the original image. Moreover, on the rest of the subregion b, our method seems to achieve higher destriping. (D) Comparison of striping intensity on subregion b: indeed, our method achieves lower striping intensity.

## 7. Discussion

VisiumHD slides exhibit slide-specific, non-periodic striping artifacts that distort bin totals and can bias downstream analyses. We presented a statistical destriping method that leverages the homogeneity of counts within nuclei to fit multiplicative row- and column-specific stripe factors in a negative binomial GLM. On synthetic data, our approach more accurately recovers stripe factors and corrected counts than `bin2cell` and related baselines. Across four public VisiumHD slides, it consistently reduces striping while better preserving large-scale total-count structure and avoiding artifacts such as edge effects, macro-stripes, and inflated regions introduced by competing methods; supplementary analyses further indicate benefits for downstream cell typing and differential expression.

## 8. Computational requirements

Across four real datasets (61k-448k nuclei), GLM fitting required 8-50 minutes and 0.7-3.2 GB RAM; the destriping step completed in under 1 minute. Full benchmarks are in Supplementary Table S1.

## 10. Acknowledgments

This research was supported by the ETH AI Center through ETH AI Center doctoral fellowships to Paola Malsot and Malte Londschien.

## Footnotes

1 The Euclidean distance is normalized by the mean of the smoothed total-count profile of the original image multiplied by the square root of the number of lanes.

