## Supplementary Material for "Striping artifact removal in VisiumHD data through nuclear counts modeling"

### S1. Introduction

In the following supplementary material, we include additional results that were omitted from the main text for the sake of brevity and clarity. These notably include the benchmarks shown in the main text with additional baselines. Moreover, we show qualitative destriping results on the mouse embryo and mouse brain slides, where similar conclusions can be drawn as those derived from the two other datasets discussed in the main text.

### S2. Additional baselines

In addition to the baselines presented in Section 5.1, this section describes supplementary baseline methods and variants that were evaluated but omitted from the main text for conciseness.

#### S2.1. Variants of b2c-sym

**b2c-sym-nucl.**
In this variant, row-wise and column-wise quantiles are estimated using only bins corresponding to nuclei, while the resulting stripe-factors are applied to all bins.

**b2c-sym-q=0.5.**
In this variant, the $99^{\text{th}}$ quantile used in the standard **b2c-sym** procedure is replaced by the median (i.e., the $50^{\text{th}}$ quantile) when estimating row-wise and column-wise stripe-factors.

**b2c-sym-nucl-q=0.5.**
In this variant, row-wise and column-wise stripe-factors are estimated using only nuclear bins and the median ($50^{\text{th}}$ quantile) of the corresponding count distributions.

#### S2.2. Ablation variants

We consider ablations of our method for estimating $(\mathbf{c}, \mathbf{h}, \mathbf{w})$ (see Section 3 for a complete description of the optimization objective). In the optimization objective, our method uses a selective $\ell_2$ regularizer

$$\mathcal{R}(\beta) = \beta^\top D\beta,$$

where $D$ is diagonal with nonzero entries only for the components of $\beta$ corresponding to $\log(\mathbf{h})$ and $\log(\mathbf{w})$ (and zeros for $\log(\mathbf{c})$). Moreover, we start the optimization procedure by initializing $\beta$ in correspondence to $h_i = 1$, $w_j = 1$, and

$$c_p = \frac{\sum_{i,j} k_{ij}\, \mathbf{1}_{\{p(i,j)=p\}}}{\sum_{i,j} \mathbf{1}_{\{p(i,j)=p\}}}.$$

**ours_N.I.**
Default `glum` initialization is used for $\beta$ instead of our initialization, while keeping the objective and regularizer unchanged. For the synthetic data (all three seeds) and the mouse embryo data, this variant produced infinite values in the destriped output; consequently, some metrics are undefined.

**ours_P2.I.**
The selective regularizer is replaced by an isotropic $\ell_2$ penalty on the full parameter vector,

$$\mathcal{R}(\beta) = \beta^\top I\beta,$$

thereby regularizing $\log(\mathbf{c})$ in addition to $\log(\mathbf{h})$ and $\log(\mathbf{w})$. Initialization is unchanged.

#### S2.3. **Median-ratio stripe estimator (MRSE)**

We also derive a simple stripe estimator inspired by our hypothesis of homogeneous counts within individual nuclei, which we refer to as the median-ratio stripe estimator (MRSE). For each bin $(i, j)$ belonging to a nucleus, we first compute the ratio between the bin count $k_{ij}$ and the mean count of the corresponding nucleus,

$$r_{ij} = \frac{k_{ij}}{\text{mean}(\{k_{i'j'} : p(i',j') = p(i,j)\})}, \tag{S1}$$

where $p(i, j)$ denotes the nucleus to which bin $(i, j)$ is assigned. Under the homogeneity assumption, these ratios reflect systematic row-wise and column-wise stripe effects.

Stripe-factors are then estimated by taking the medians of the ratios across nuclear bins in the corresponding row or column:

$$h_i = \text{median}(\{r_{ij} : (i,j) \text{ is a nuclear bin}\}), \qquad w_j = \text{median}(\{r_{ij} : (i,j) \text{ is a nuclear bin}\}). \tag{S2}$$

Finally, corrected bin counts are obtained by dividing each bin count by the product of its corresponding row-wise and column-wise stripe-factors,

$$k_{ij}^{\text{corr}} = \frac{k_{ij}}{h_i\, w_j}. \tag{S3}$$

### S3. Supplementary Figures

#### S3.1. Results on synthetic data

On the synthetic data benchmark, we see that b2c-derived baseline b2c-sym-nucl achieves a slightly lower error than b2c, but still considerably higher than ours (see Fig. S4). Interestingly, MRSE achieves a much lower error than b2c-derived baselines (see Fig. S4) and also a better preservation of global structure and lower striping intensity (see Fig. S1). On Figure S1, we also observe that our simulation benchmark recapitulates observations from real data, notably that b2c-derived baselines introduce considerable global structure alteration and that our method is more effective at reducing the striping intensity.

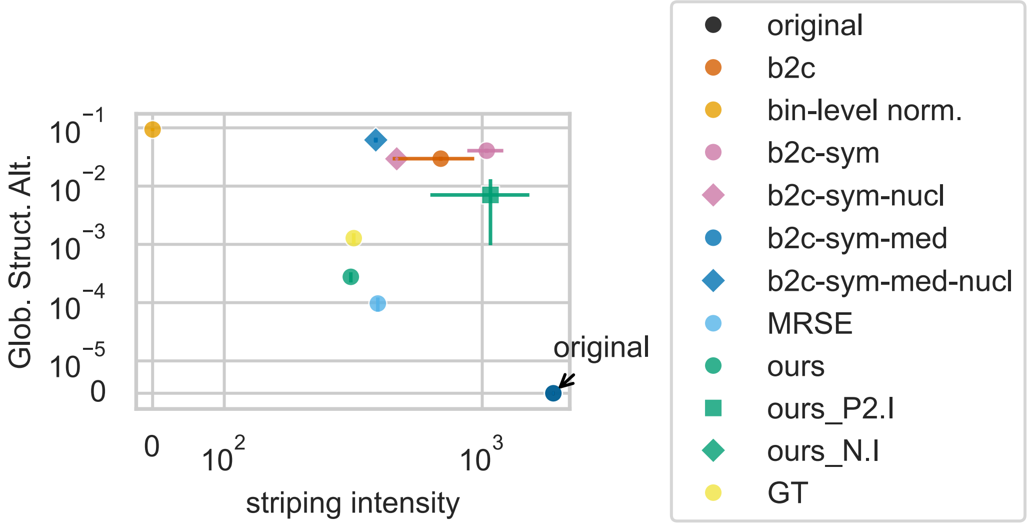


Fig. S1: Tradeoff between striping-intensity reduction and global-structure preservation on synthetic data. Results are qualitatively similar to those in real data (see Fig. 3). b2c-derived baselines reduce striping intensity but substantially alter global structure. Our method achieves lower striping intensity while minimizing global-structure changes, comparable to corrected counts using ground-truth stripe factors (GT). MRSE also achieves a good trade-off, although it destripes the data less than ours. As a reference, bin-level normalization yields zero striping intensity by equalizing counts across bins. The ablation variants of our model (ours_N.I. and ours_P2.I) do not perform well. The ours_N.I. method produced destriped data containing infinite values; therefore, the corresponding metrics are undefined and are not shown in the plot. Error bars indicate the standard deviation across three data-generation seeds.

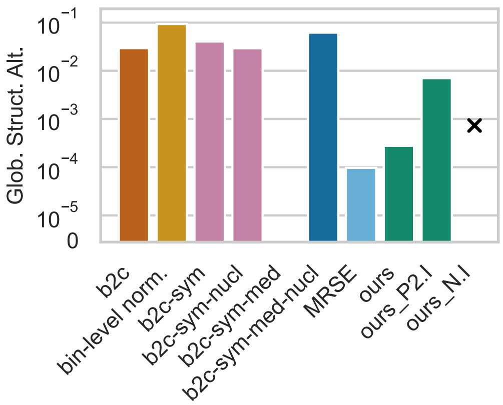


Fig. S2: Global structure alteration for different baseline variants. The ours_N.I. method produced destriped data containing infinite values; therefore, the global structure alteration score is undefined and not shown in the plot.

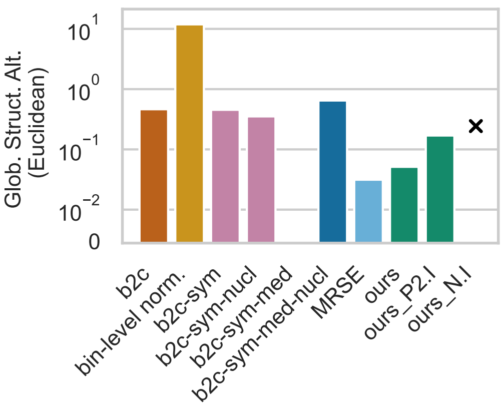


Fig. S3: Global structure alteration for different baseline variants measured with the normalized euclidean distance. The ours_N.I. method produced destriped data containing infinite values; therefore, the global structure alteration score is undefined and not shown in the plot.

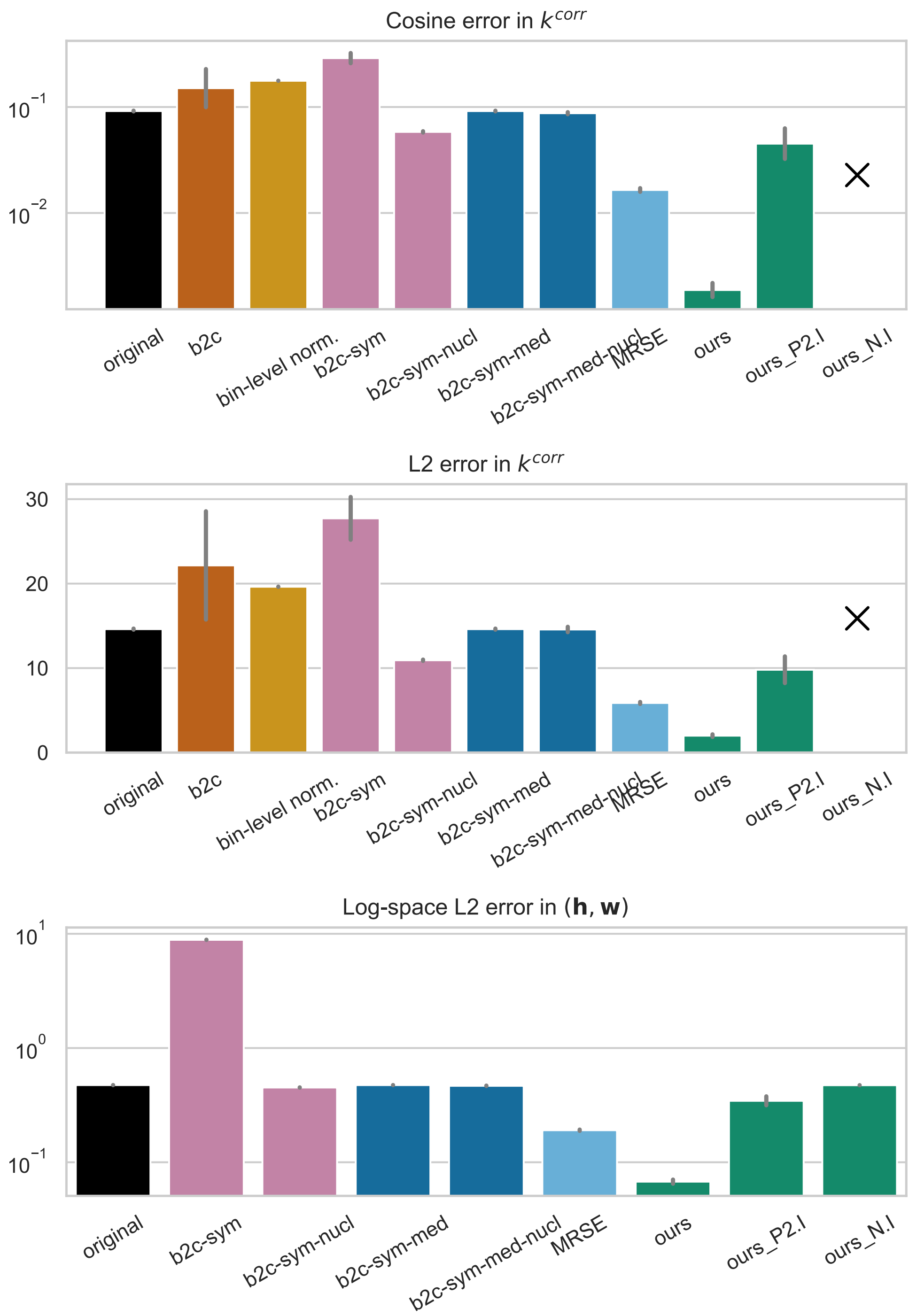


Fig. S4: Comparison of stripe-correction methods on synthetic data. Performance is evaluated using: (top) the cosine error of the corrected counts, defined as the cosine distance to the ground-truth reference shown in Fig. 2B; (middle) the $\ell_2$ version of this error, normalized by the square root of the number of bins; and (bottom) stripe-factor estimation accuracy, measured by the $\ell_2$ error between the estimated stripe factors $(\mathbf{h}, \mathbf{w})$ and the ground truth in log-space. Method b2c is omitted from the bottom panel because it does not explicitly model stripe factors. Since the method ours_N.I. generated destriped data containing infinite values, the corresponding errors in $k^{\text{corr}}$ are not computable (indicated with a cross). For reference, original denotes $\mathbf{h} = \mathbf{1}$ and $\mathbf{w} = \mathbf{1}$. Overall, our model achieves superior performance in both count correction (top, middle) and stripe-factor estimation (bottom). Error bars indicate the standard deviation across three data-generation seeds.

#### S3.2. Results on the mouse brain data

On the mouse brain data, we highlight in Figures S6 and S7 a horizontal macro-stripe artifact which is introduced by the b2c-derived baselines. Moreover, we see in Figure S9, that at the bottom edge of the slide, b2c-derived baselines produce artificially high count values, which appear as a band of higher intensity. We also show on Figure S8 a central region of the brain. There, visually, both b2c-derived methods and ours destripe the data efficiently. On this dataset, we also observed that the MRSE baseline achieves a lower global structure alteration score than our method, but achieves higher striping than our method and than b2c-derived baselines (see Fig. S14). We also see that both our ablations ours_N.I. and ours_P2.I fail to reduce the striping intensity, and the latter also introduces considerable global structure alteration.

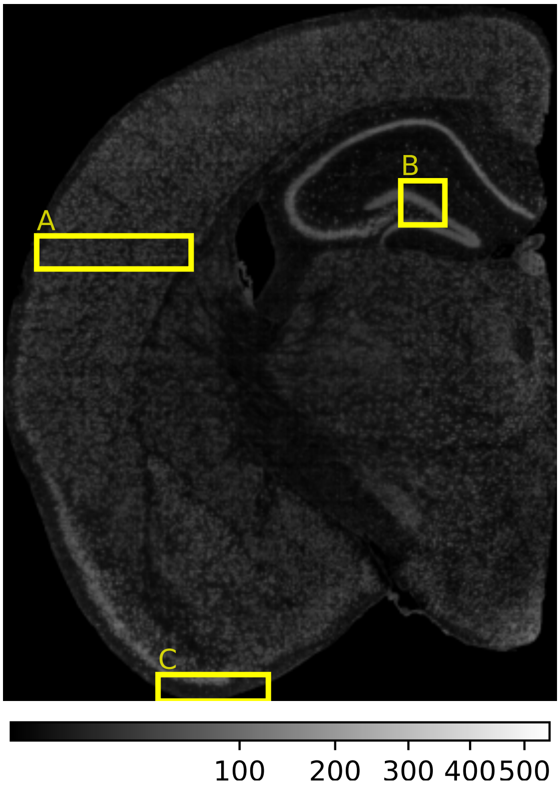


Fig. S5: Overview of VisiumHD mouse brain slide. Rectangles mark the regions shown in Figures S6 (A), S8 (B) and S9 (C).

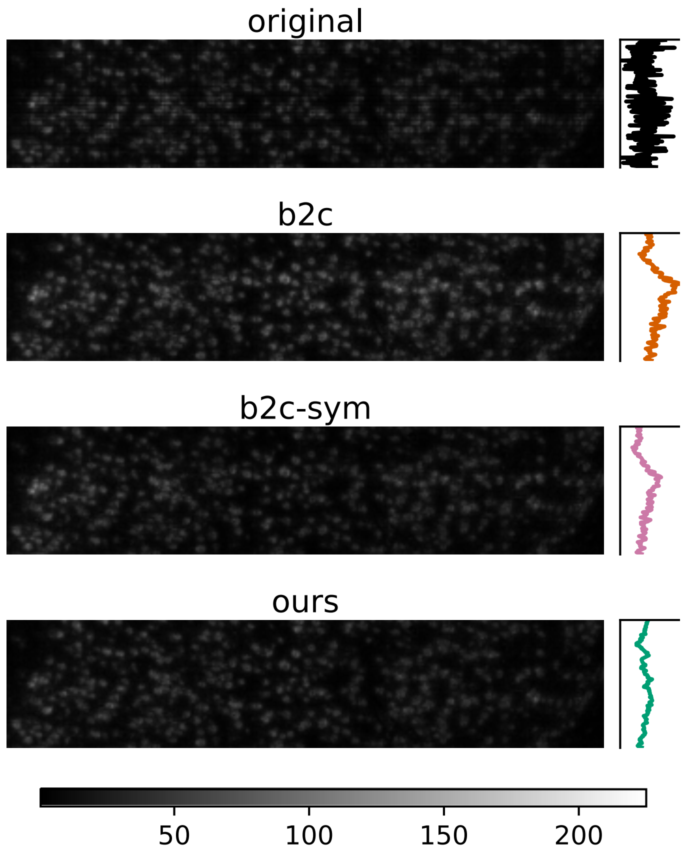


Fig. S6: Region A of mouse brain slide (see Fig. S5). In b2c and slightly in b2c-sym, a macro-stripe with higher intensity is artificially created. This artifact is not present in our method.

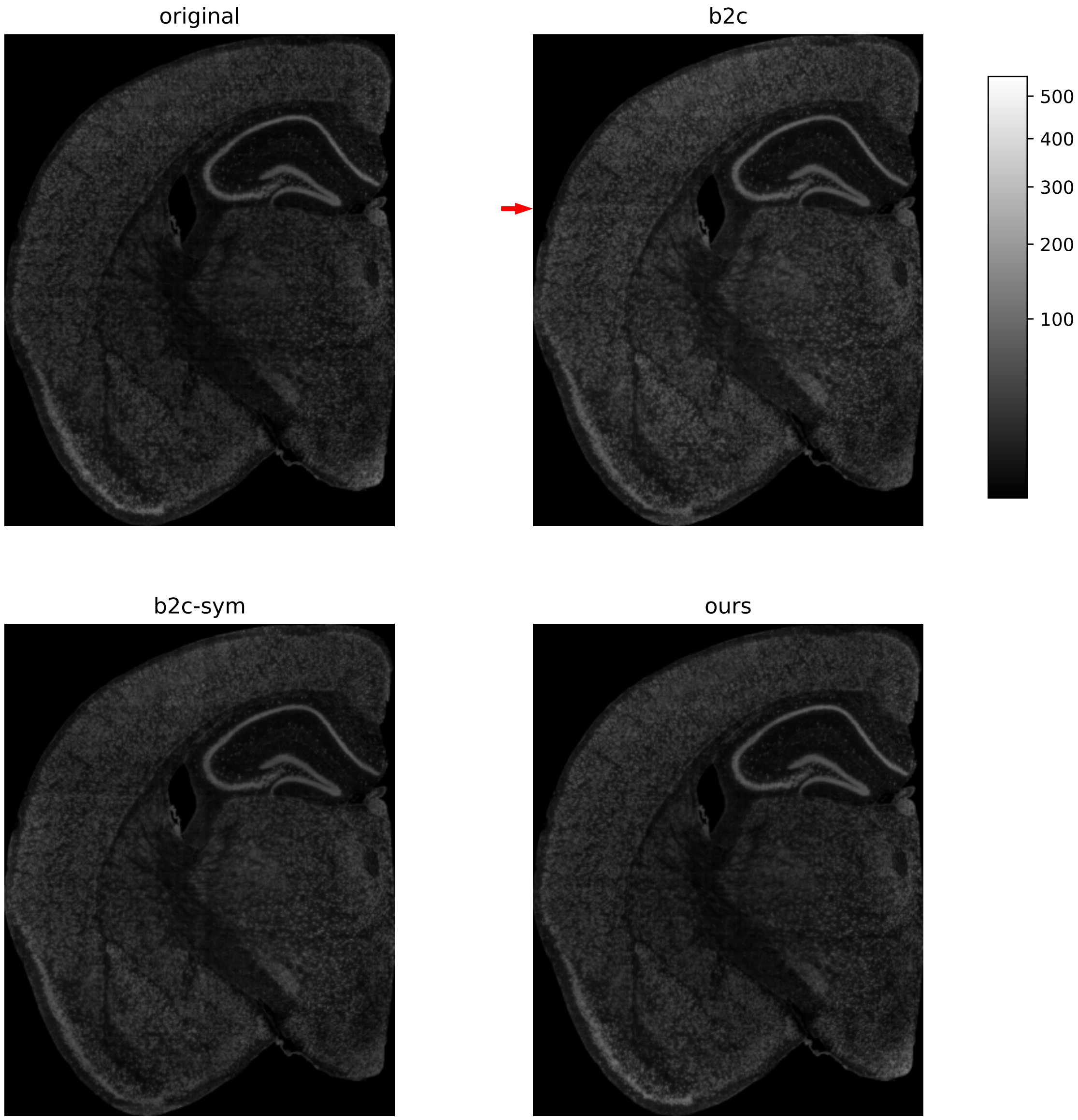


Fig. S7: Mouse brain slide. The macro-stripe shown in supplementary Fig. S6 is indicated with an arrow.

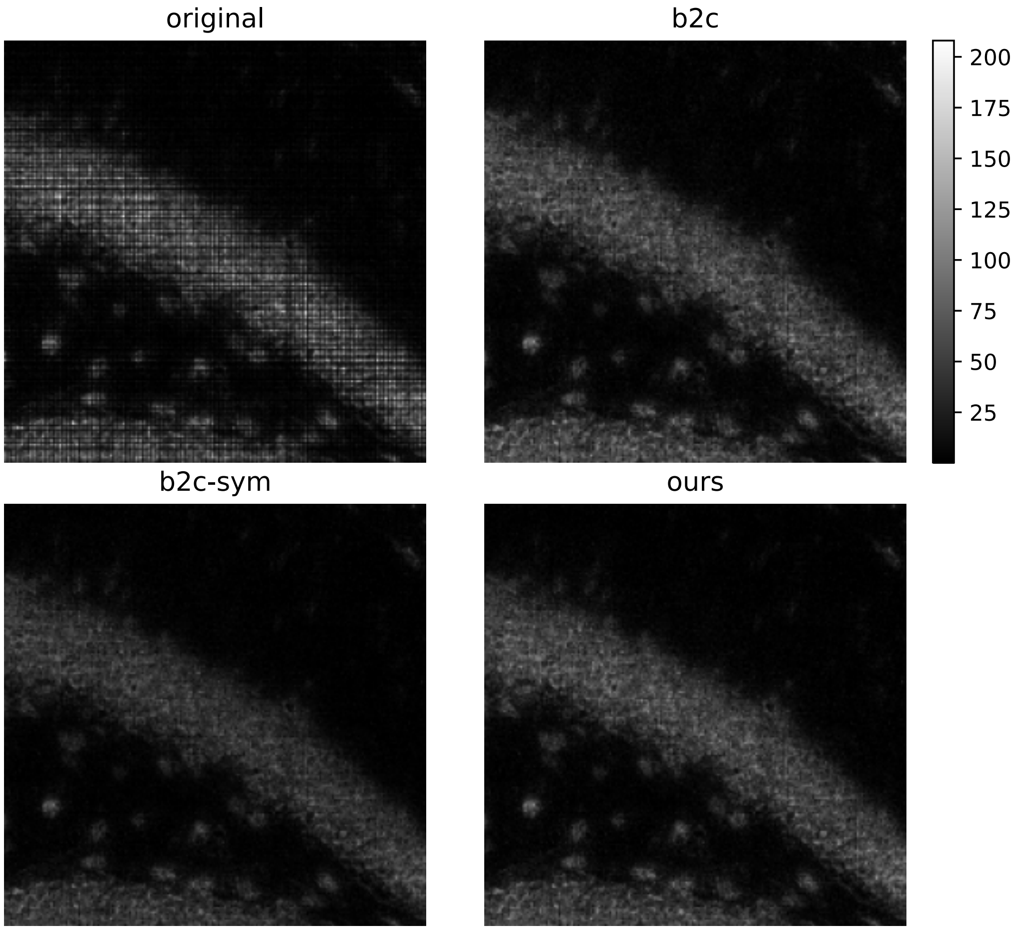


Fig. S8: Region B of mouse brain slide (see Fig. S5). This is a typical example of destriping on a zoomed region at the center of the slide.

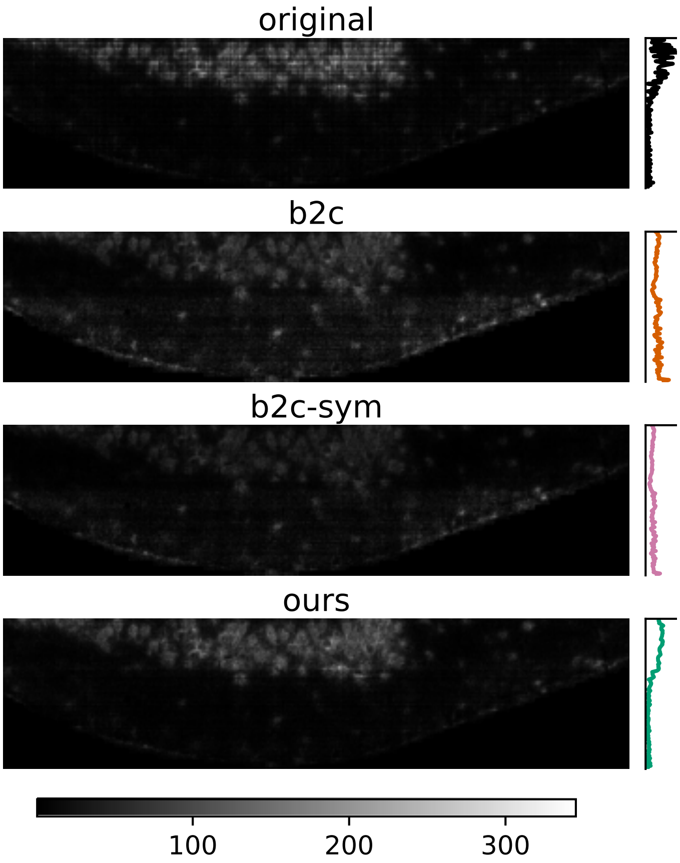


Fig. S9: Region C of mouse brain slide (see Fig. S5). In b2c and slightly in b2c-sym, horizontal stripes with higher intensity are artificially created at the bottom. This artifact is not present in our method. The plots on the right display the mean-count per lane across bins belonging to the dataset (lower right and lower left corners are empty): it is clear that b2c and b2c-sym create artificially high counts at the bottom edge.

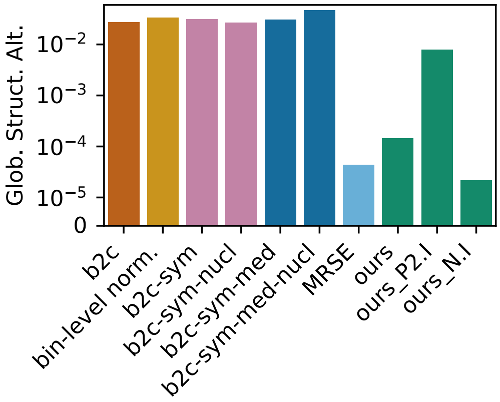


Fig. S10: Global structure alteration for different baseline variants.

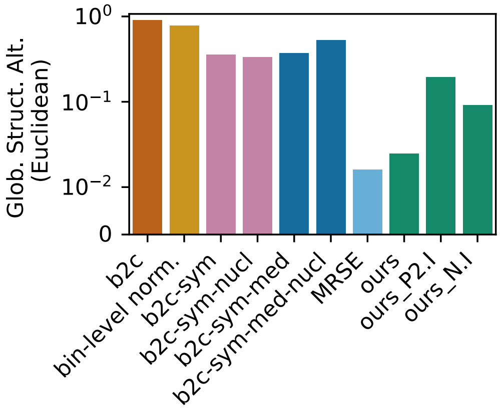


Fig. S11: Global structure alteration for different baseline variants measured with the normalized euclidean distance.

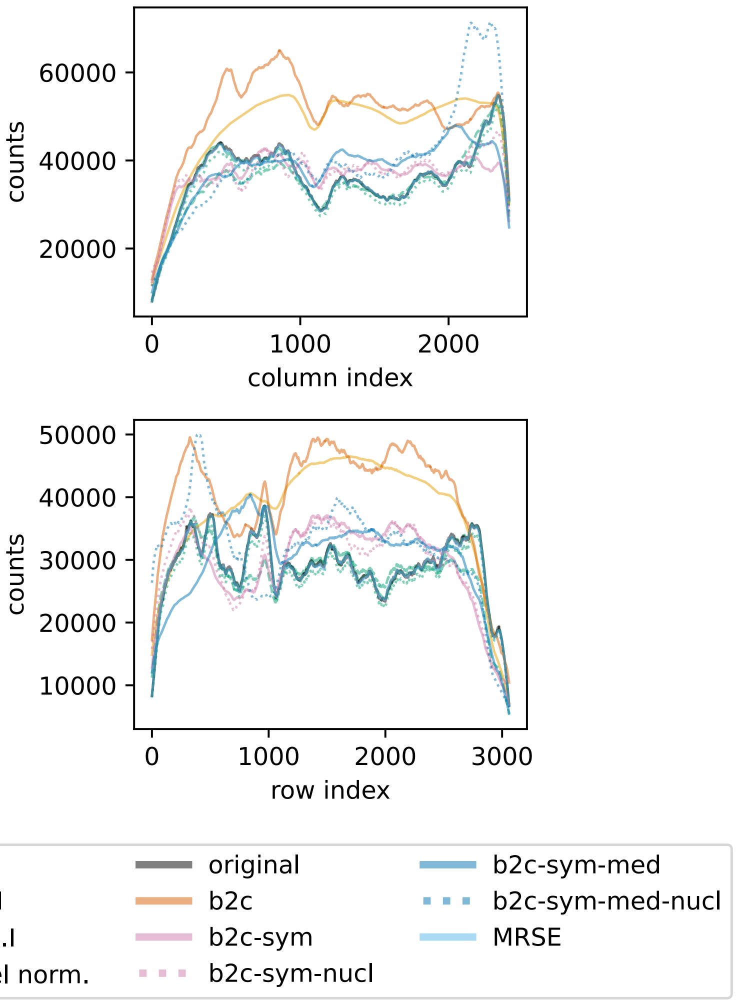


Fig. S12: Global structure profile of corrected counts for different methods. The profile is calculated by summing counts for each column (top) or row (bottom) and smoothing the resulting curve.

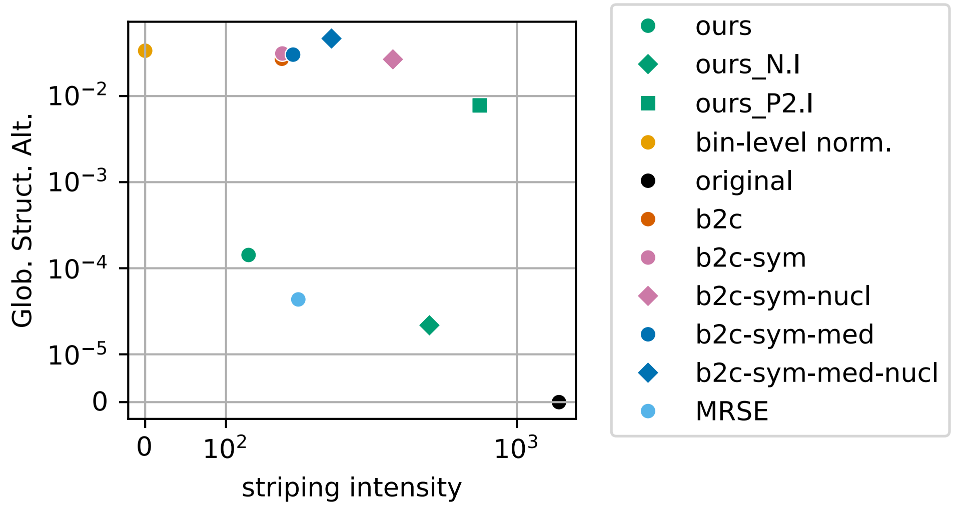


Fig. S13: Tradeoff between cytoplasmic striping-intensity reduction and global-structure preservation.

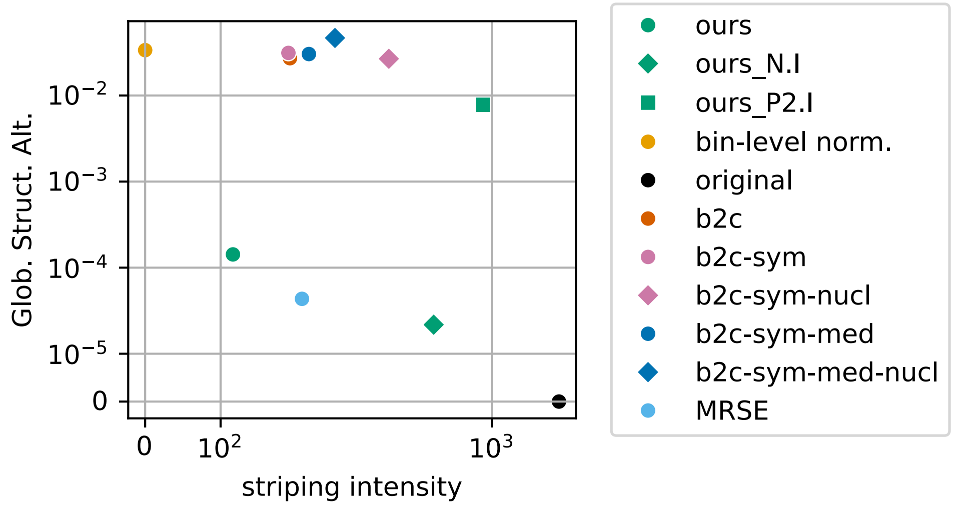


Fig. S14: Tradeoff between striping-intensity reduction and global-structure preservation. Note that here the striping intensity is not calculated only on the cytoplasm as in main figure, but on the whole total counts image including the nuclei.

#### S3.3. Results on the mouse embryo data

The mouse embryo slide gives another example where b2c-derived baselines introduce large regions of artificially high count values (see Fig. S15 and S16). Moreover, we show a portion of the slide where visually (and quantitatively) our method removes the striping artifact better than others (see Fig. S17 and S18). As before, the baseline MRSE alters less the global count structure than our method, but at the cost of lower striping intensity reduction. Note that the ours_N.I. ablation variant produced infinite values in the destriped data and is therefore missing from Figures S19, S20, S21, S22 and S23.

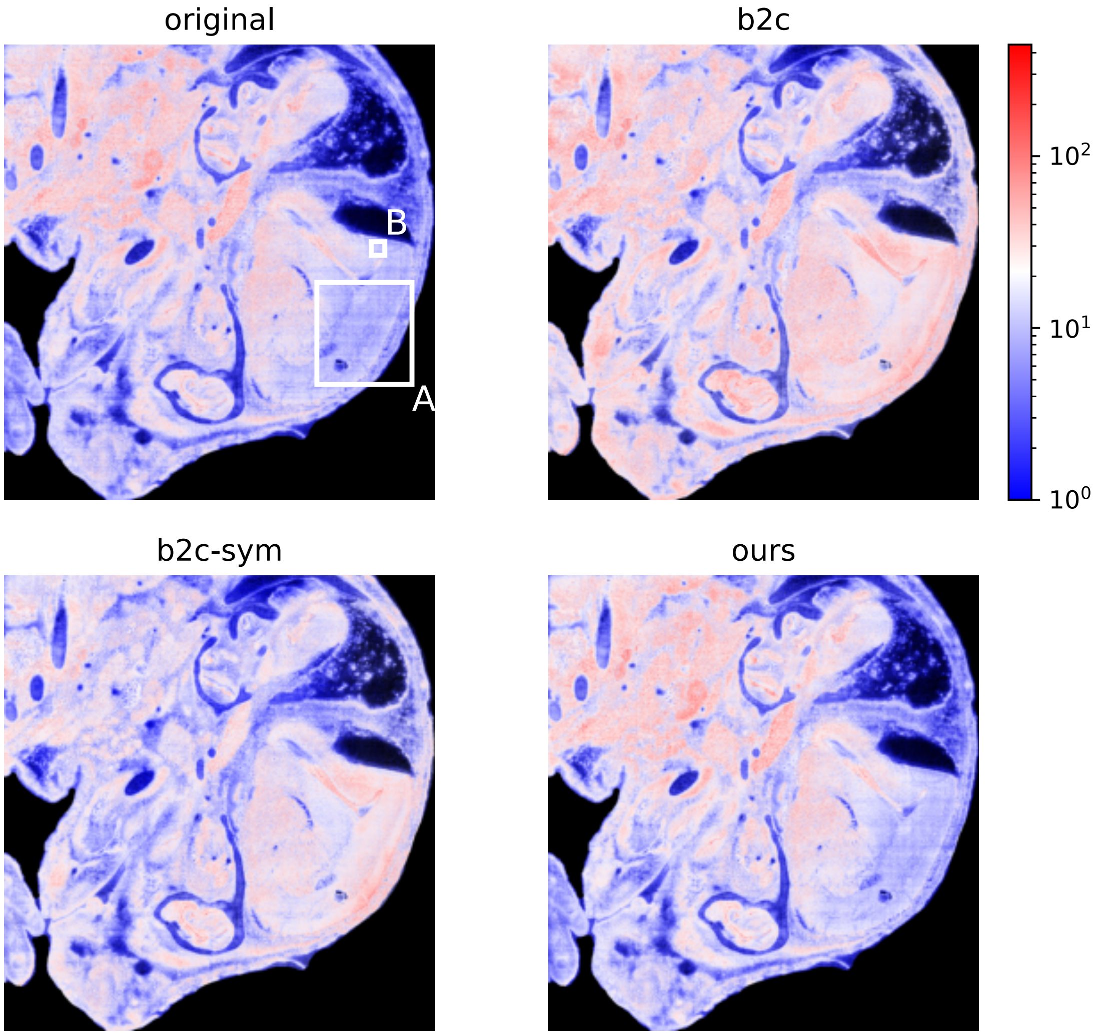


Fig. S15: Total count image of the whole mouse embryo slide for b2c, b2c-sym and our method. b2c and b2c-sym considerably alter the global count image, especially on the lower-right region (A), where artificially high counts are created. Our method does not exhibit this artifact.

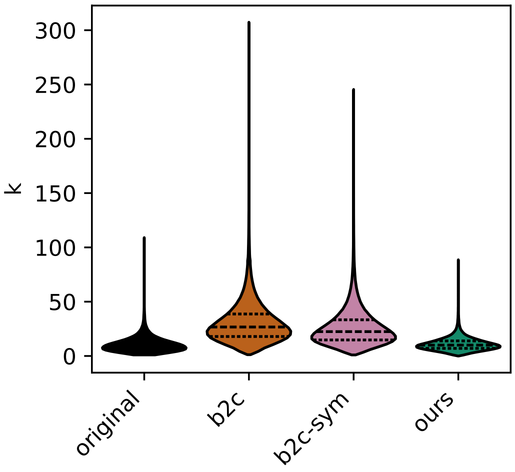


Fig. S16: Count distribution in the subregion (A) depicted in Figure S15. b2c and b2c-sym corrected count data display considerably higher values than in the original data.

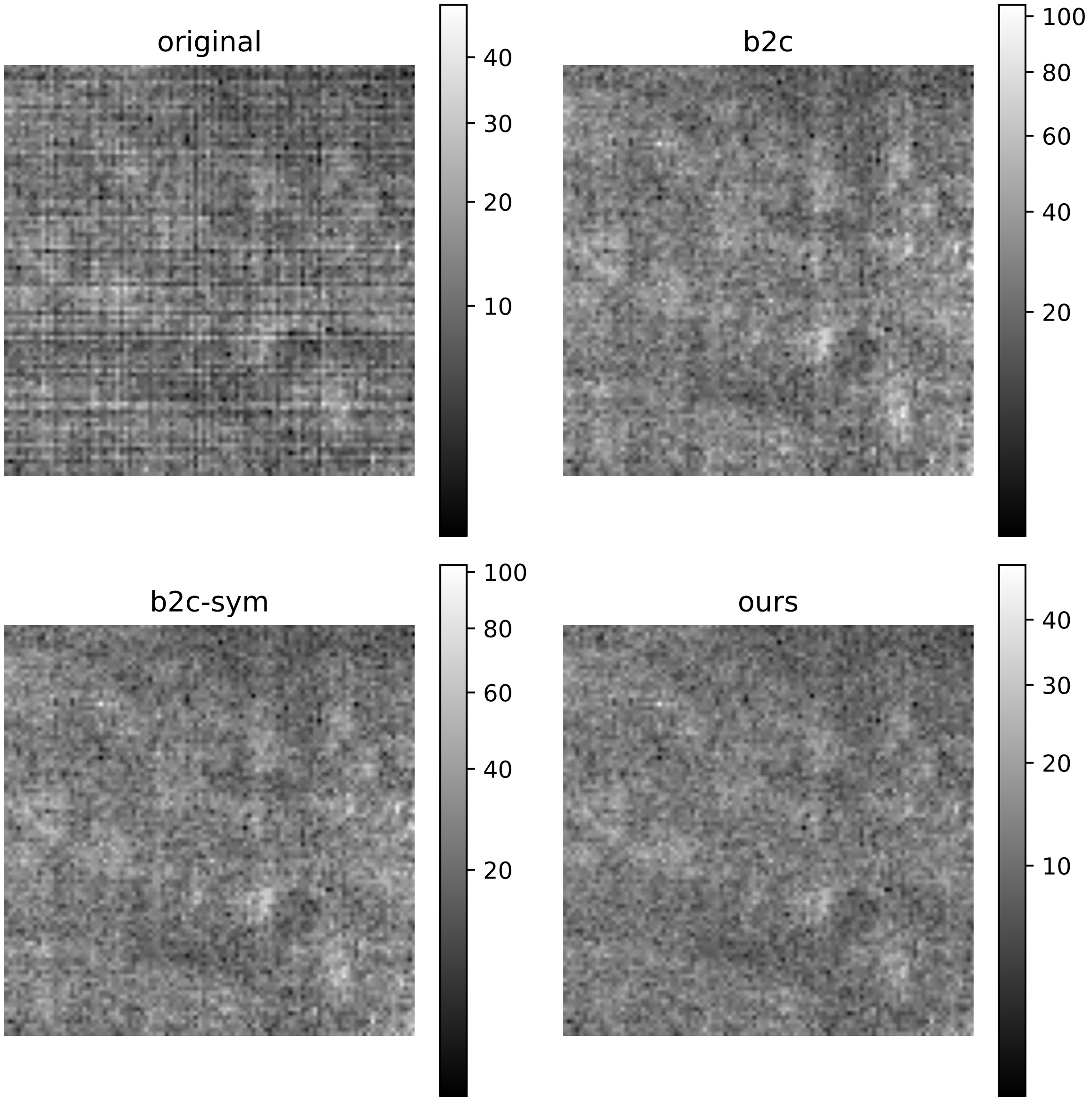


Fig. S17: Corrected total counts image for the subregion (B) depicted in Figure S15. Visually, it seems that our method removes the striping artifact slightly better. This is further confirmed in Figure S18.

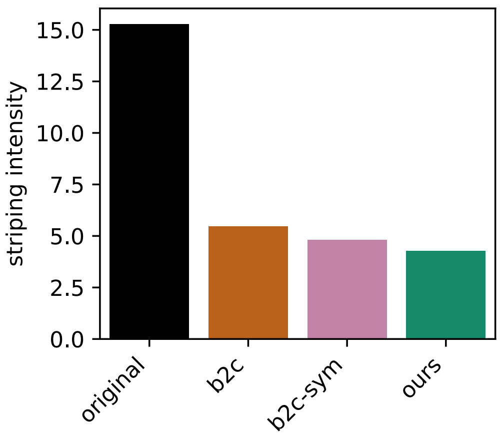


Fig. S18: Cytoplasmic striping intensity on the subregion (B) depicted in Figure S15. Our method achieves lower striping intensity than other methods.

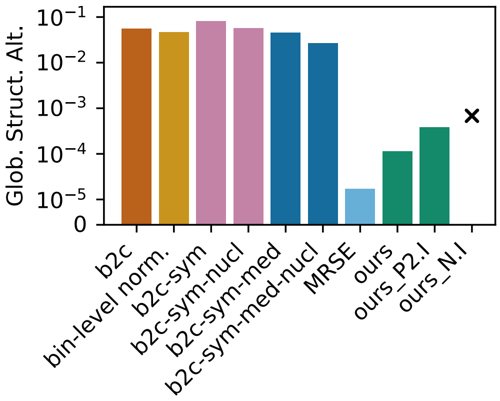


Fig. S19: Global structure alteration for different baseline variants.

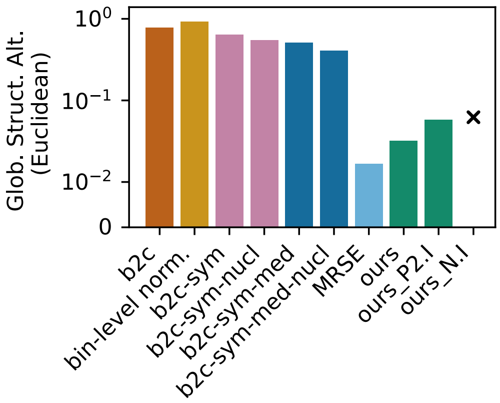


Fig. S20: Global structure alteration for different baseline variants measured with the normalized euclidean distance.

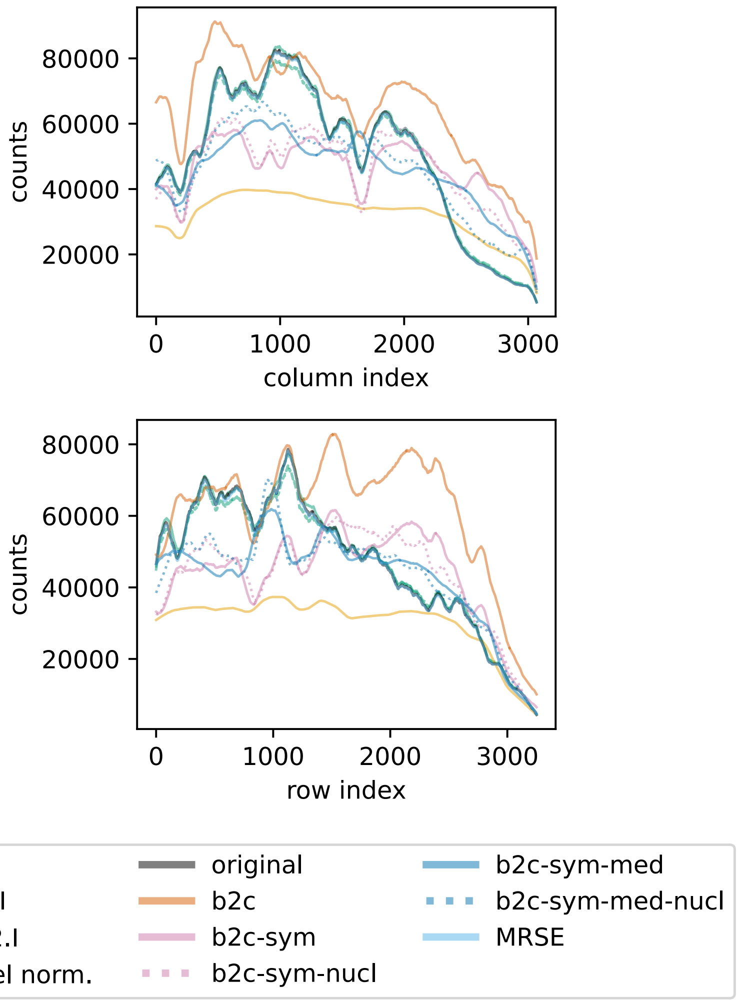


Fig. S21: Global structure profile of corrected counts for different methods. The profile is calculated by summing counts for each column (top) or row (bottom) and smoothing the resulting curve.

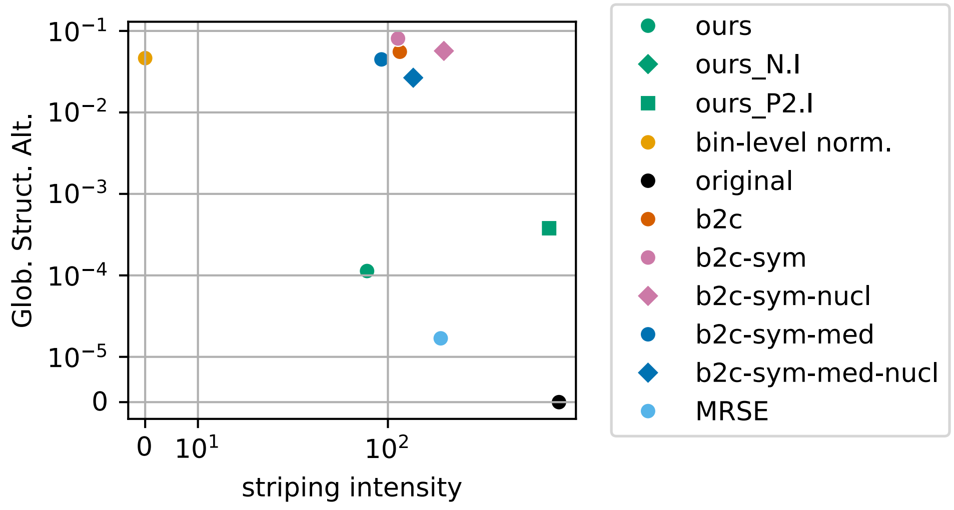


Fig. S22: Tradeoff between cytoplasmic striping-intensity reduction and global-structure preservation.

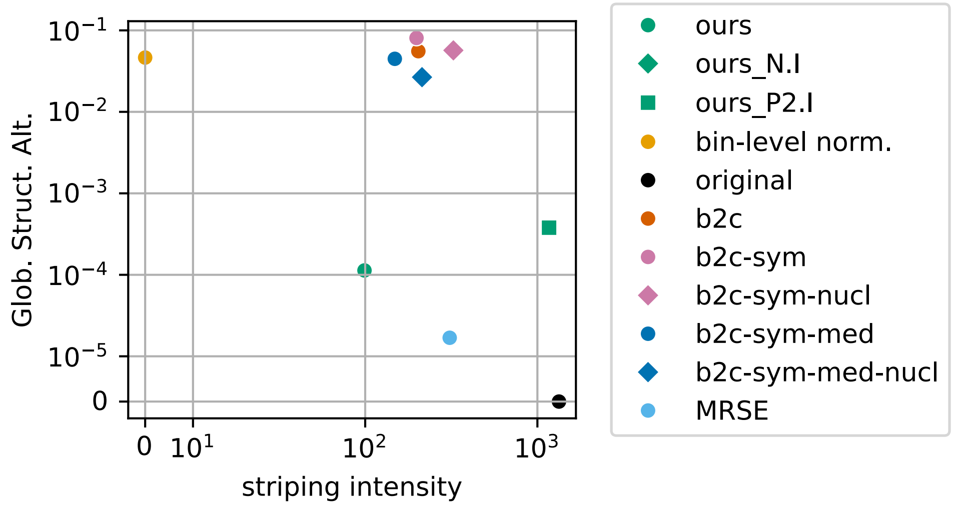


Fig. S23: Tradeoff between striping-intensity reduction and global-structure preservation. Note that here the striping intensity is not calculated only on the cytoplasm as in main figure, but on the whole total counts image including the nuclei.

#### S3.4. Results on the zebrafish head data

On the zebrafish dataset, our method achieves a slightly higher (but comparable) cytoplasmic striping intensity than b2c, while introducing considerably lower global structure alteration. Also note that MRSE totally failed to destripe the image, showing that this simple estimator produces unstable solutions.

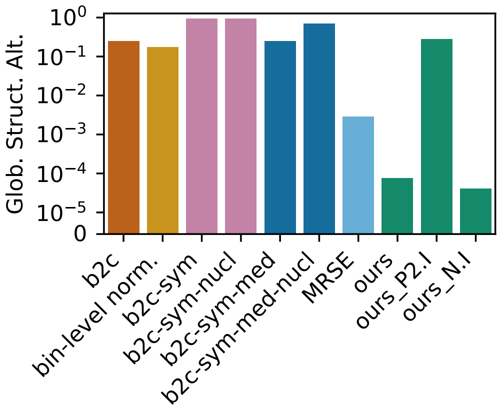


Fig. S24: Global structure alteration for different baseline variants.

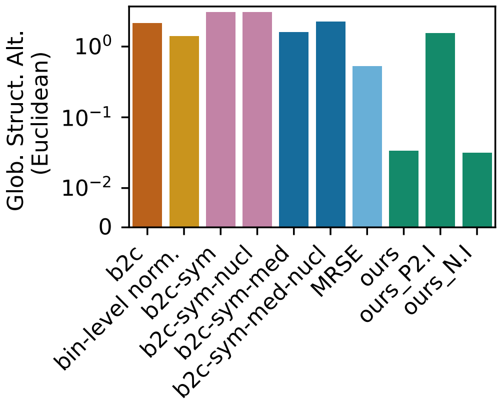


Fig. S25: Global structure alteration for different baseline variants measured with the normalized euclidean distance.

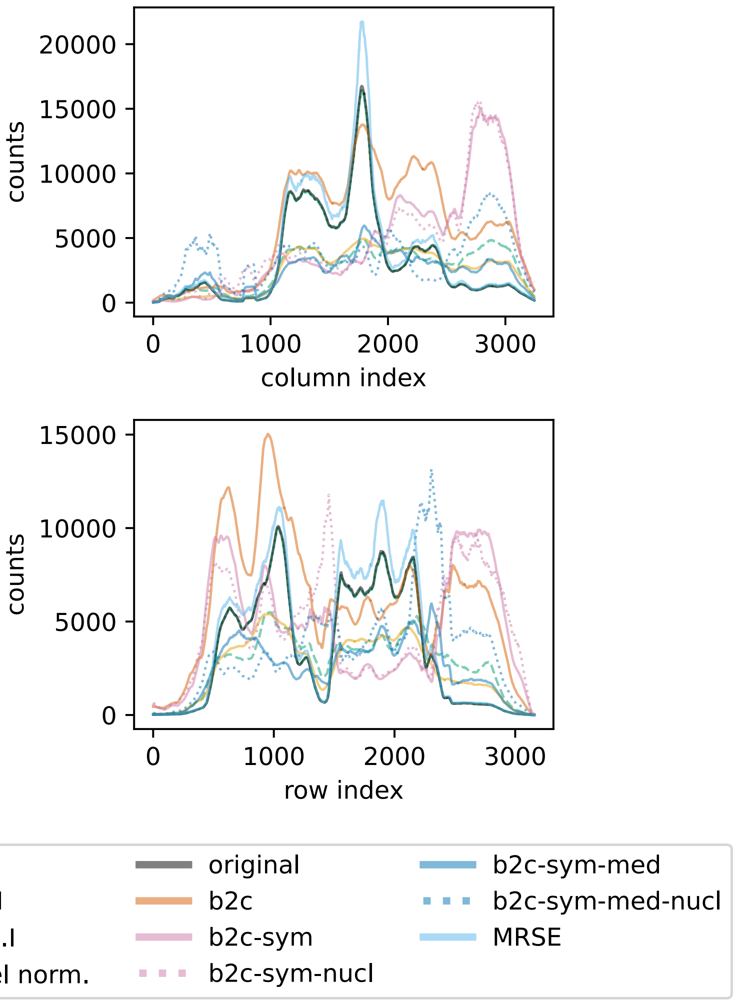


Fig. S26: Global structure profile of corrected counts for different methods. The profile is calculated by summing counts for each column (top) or row (bottom) and smoothing the resulting curve.

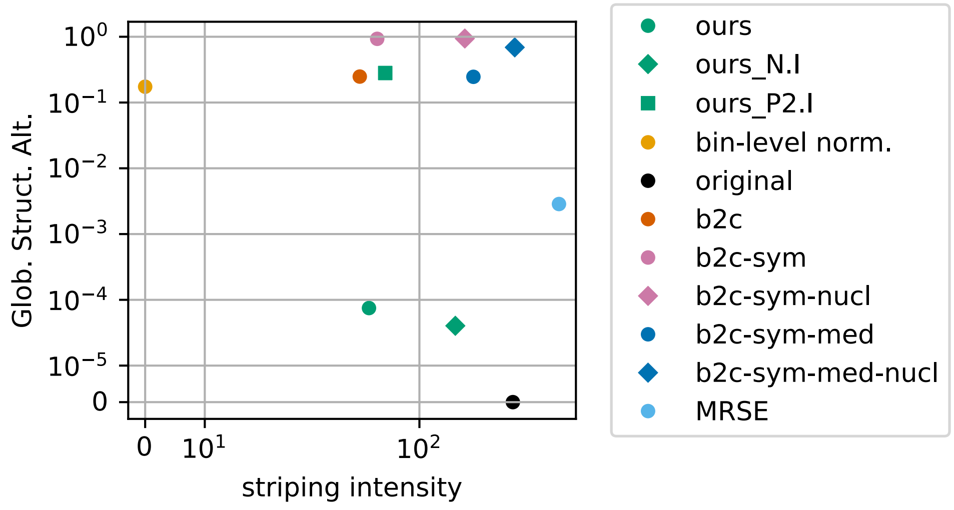


Fig. S27: Tradeoff between cytoplasmic striping-intensity reduction and global-structure preservation.

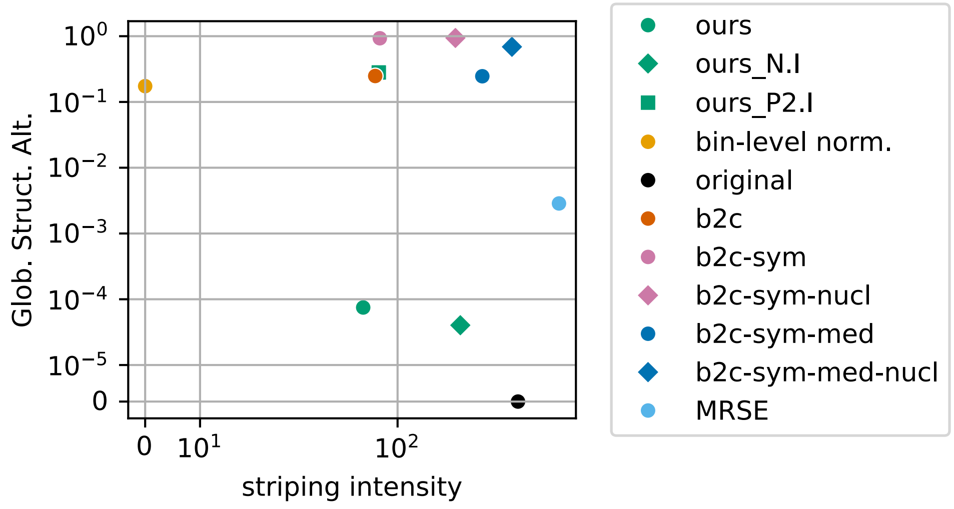


Fig. S28: Tradeoff between striping-intensity reduction and global-structure preservation. Note that here the striping intensity is not calculated only on the cytoplasm as in main figure, but on the whole total counts image including the nuclei.

S3.5. Results on the human lymph node data

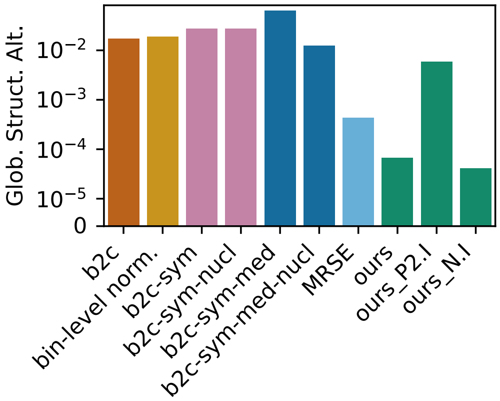


Fig. S29: Global structure alteration for different baseline variants.

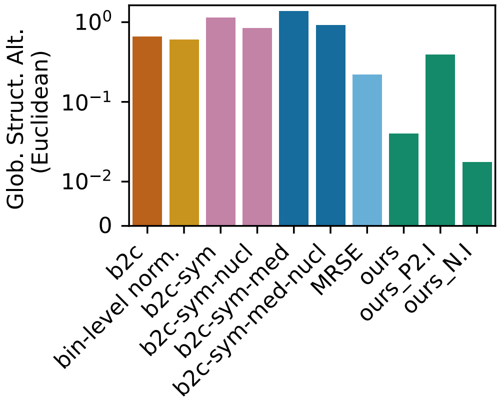


Fig. S30: Global structure alteration for different baseline variants measured with the normalized euclidean distance.

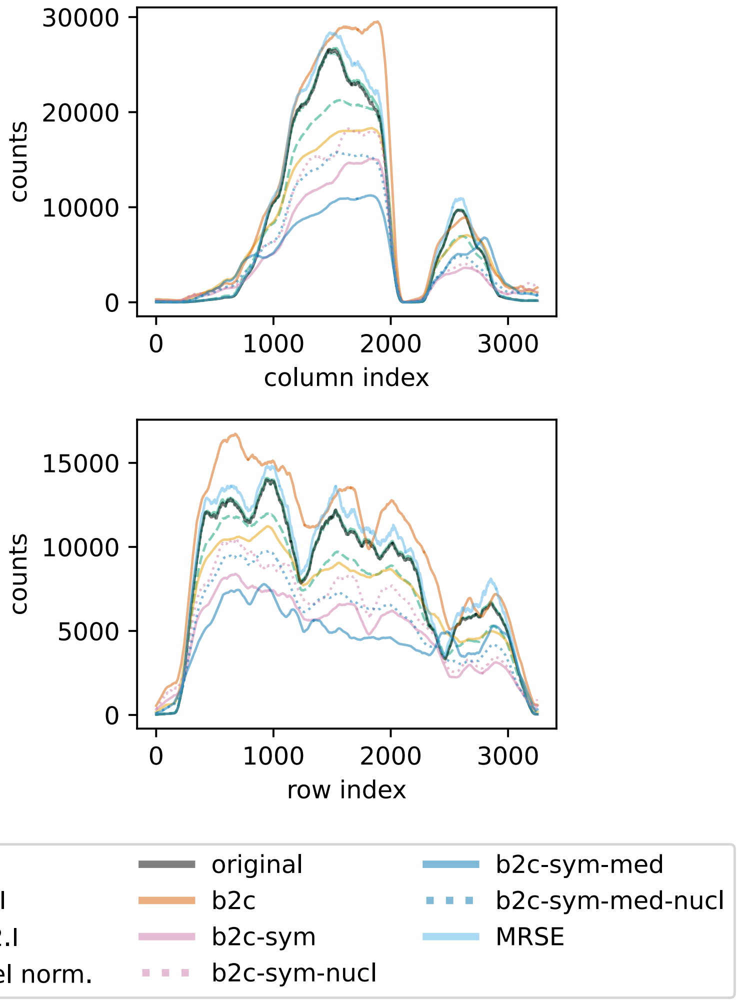


Fig. S31: Global structure profile of corrected counts for different methods. The profile is calculated by summing counts for each column (top) or row (bottom) and smoothing the resulting curve.

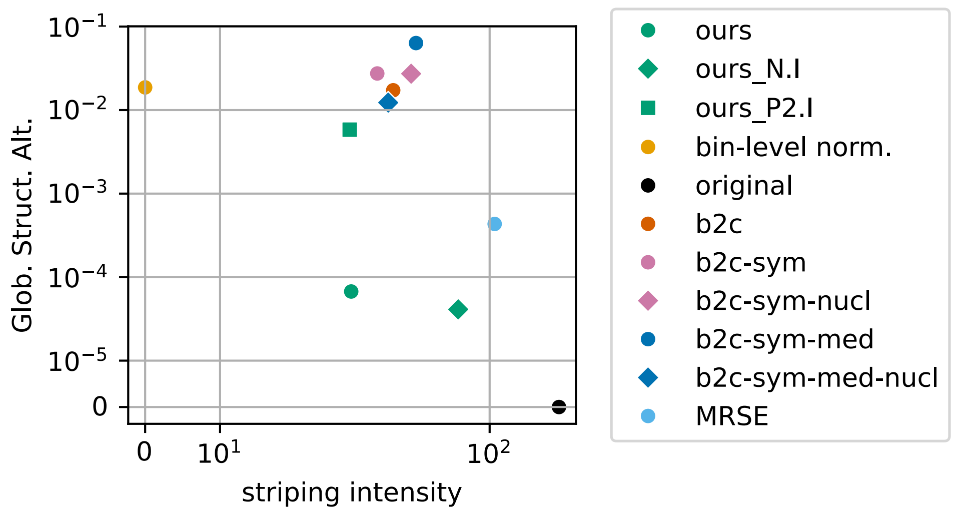


Fig. S32: Tradeoff between cytoplasmic striping-intensity reduction and global-structure preservation.

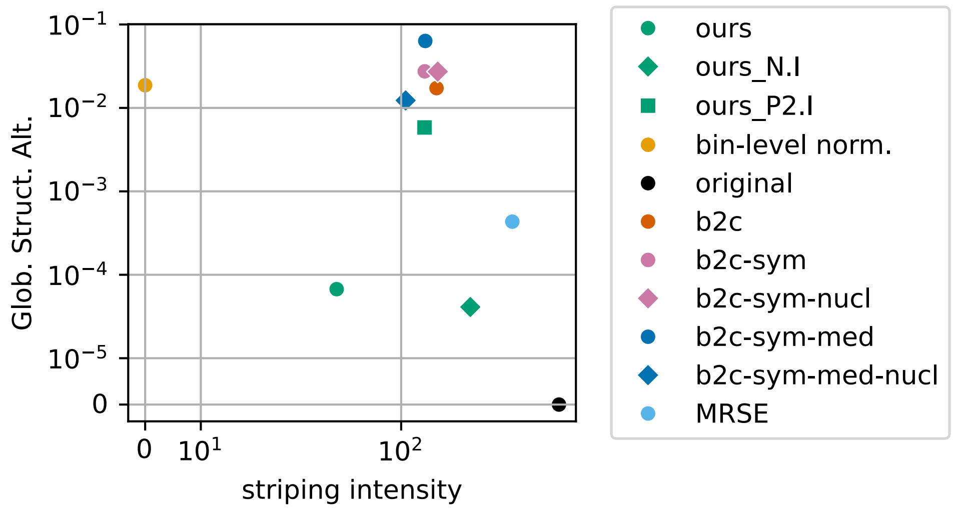


Fig. S33: Tradeoff between striping-intensity reduction and global-structure preservation. Note that here the striping intensity is not calculated only on the cytoplasm as in main figure, but on the whole total counts image including the nuclei.

### S4. Robustness to segmentation accuracy

In this section, we evaluate the robustness of our method to segmentation errors. We consider three segmentation failure modes: missing cells, undersegmentation, and oversegmentation. We assess their impact on both one synthetic mouse brain dataset and on the human lymph node dataset. We selected the latter because it contains regions of densely packed cells, which are expected to be particularly sensitive to oversegmentation.

We introduce errors into the segmentation mask as follows:

**Missing cells.** We keep $p\%$ of nuclei from the segmentation mask.

**Oversegmentation.** We split $p\%$ of nuclei into two *child* nuclei.

**Undersegmentation.** We construct a graph in which nuclei are connected by an edge when they are adjacent in the segmentation mask. We then simulate undersegmentation by merging nuclei along $p\%$ of these edges. Note that nuclei do not all have the same number of neighbors; therefore, the effect of randomly merging cells is not uniform. This effect is illustrated in Figures S35 and S34. To assess the behavior of the method when all nuclei are assigned the same label, irrespective of their degree of connectivity, we introduce an additional baseline, **ours (collapsed label)**.

In each experiment, nuclei or edges were selected at random. For increasing values of $p$, selections were performed in a nested manner, and each experiment was repeated with three random seeds.

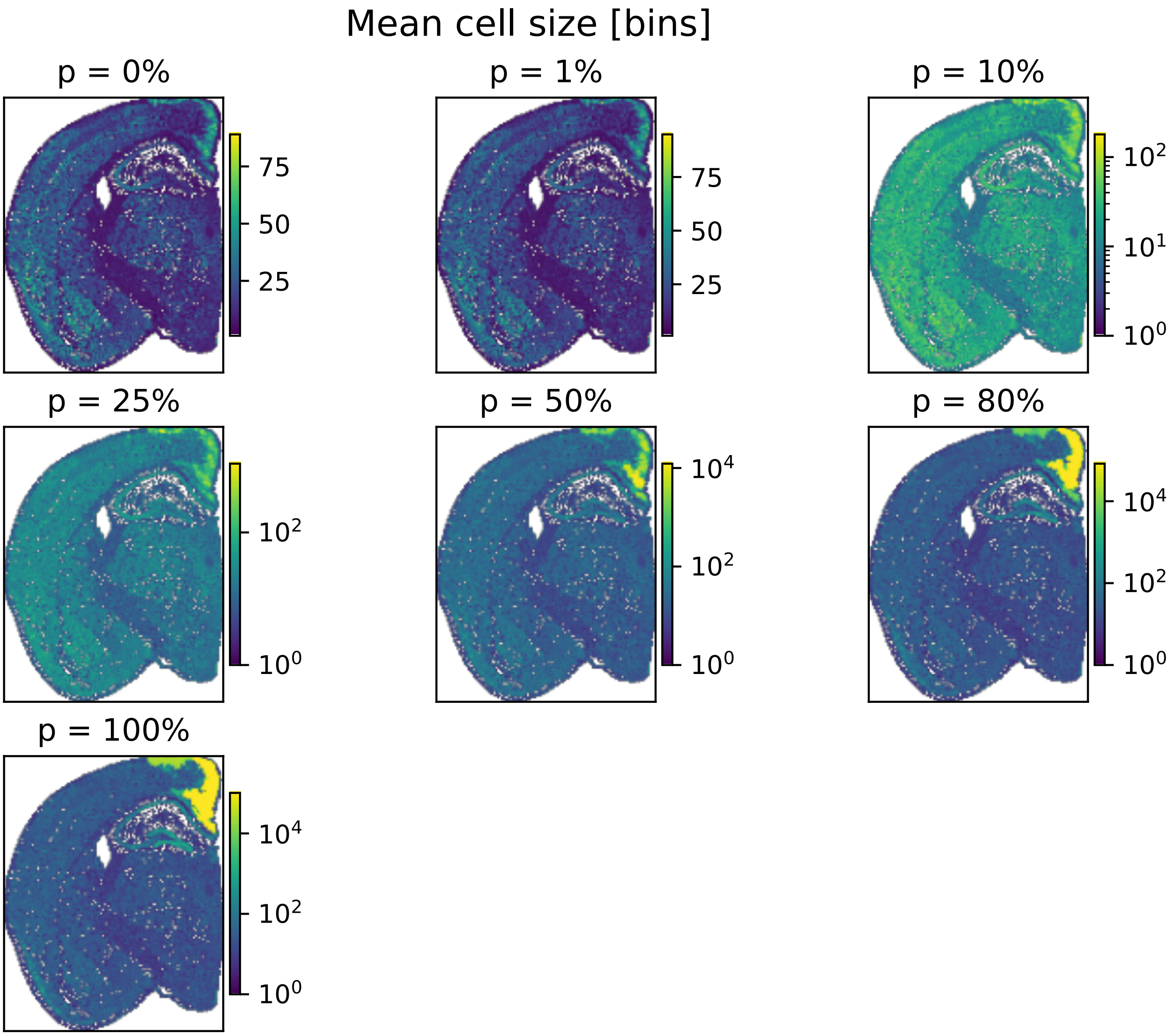


Fig. S34: Average cell sizes across merging rates (for one seed) on the synthetic mouse brain dataset.

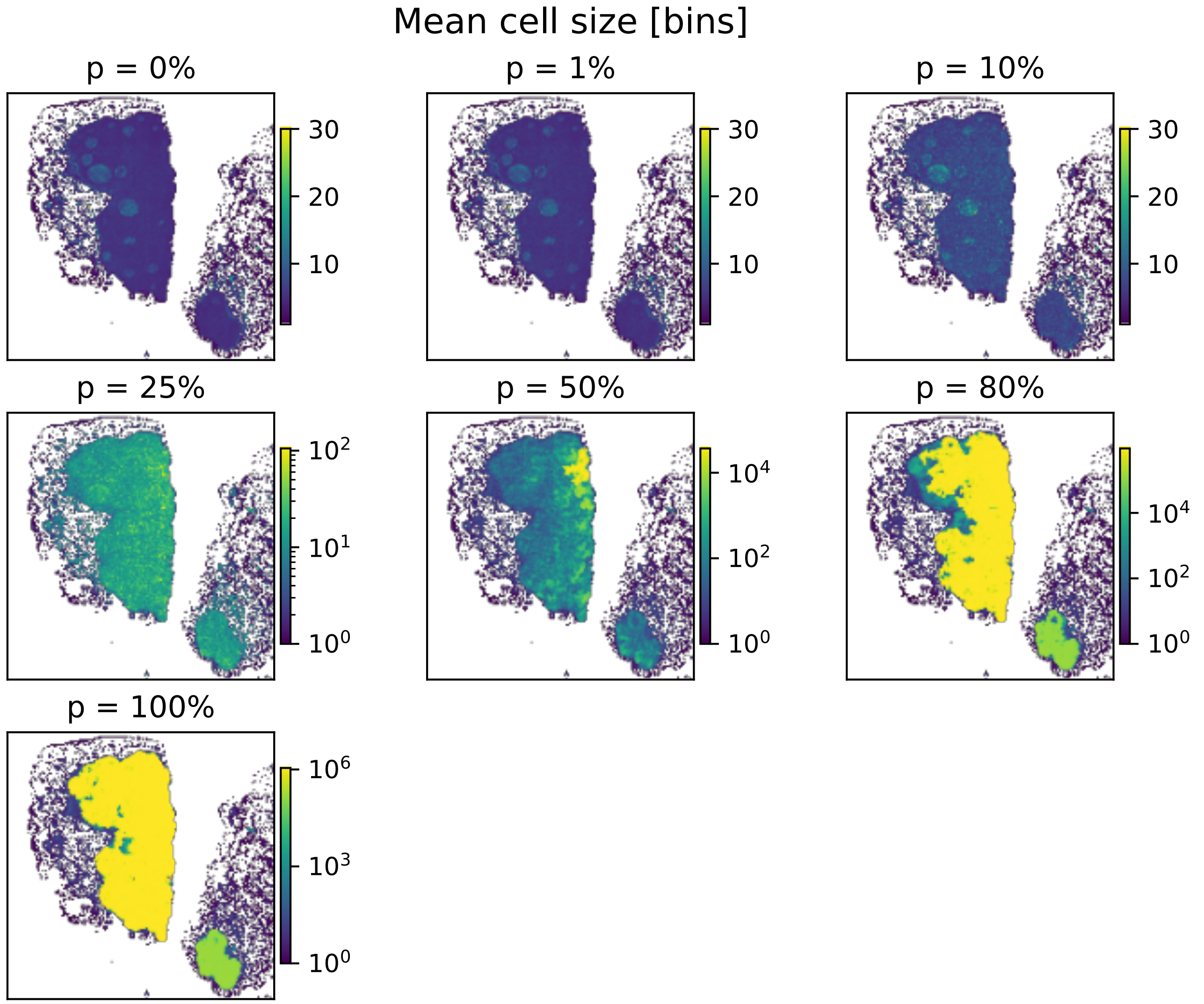


Fig. S35: Average cell sizes across merging rates (for one seed) on the human lymph node dataset.

### S4.1. Results

#### S4.1.1. Missing cells

Our method is highly robust to missing cells in both the synthetic mouse brain and human lymph node datasets (Figures S36, S37 and S38). On the synthetic brain data, performance remains satisfactory down to 10% of retained cells, as measured by the error in corrected counts, stripe factor estimation accuracy, striping intensity reduction, and global structure alteration score. On the human lymph node dataset, satisfactory trade-offs between global structure alteration and striping intensity are maintained down to 50% of retained cells. As the number of observed cells decreases, fewer observations are available to estimate each stripe factor, and the solution progressively approaches the original one, i.e. $(\mathbf{h} = \mathbf{1}, \mathbf{w} = \mathbf{1})$.

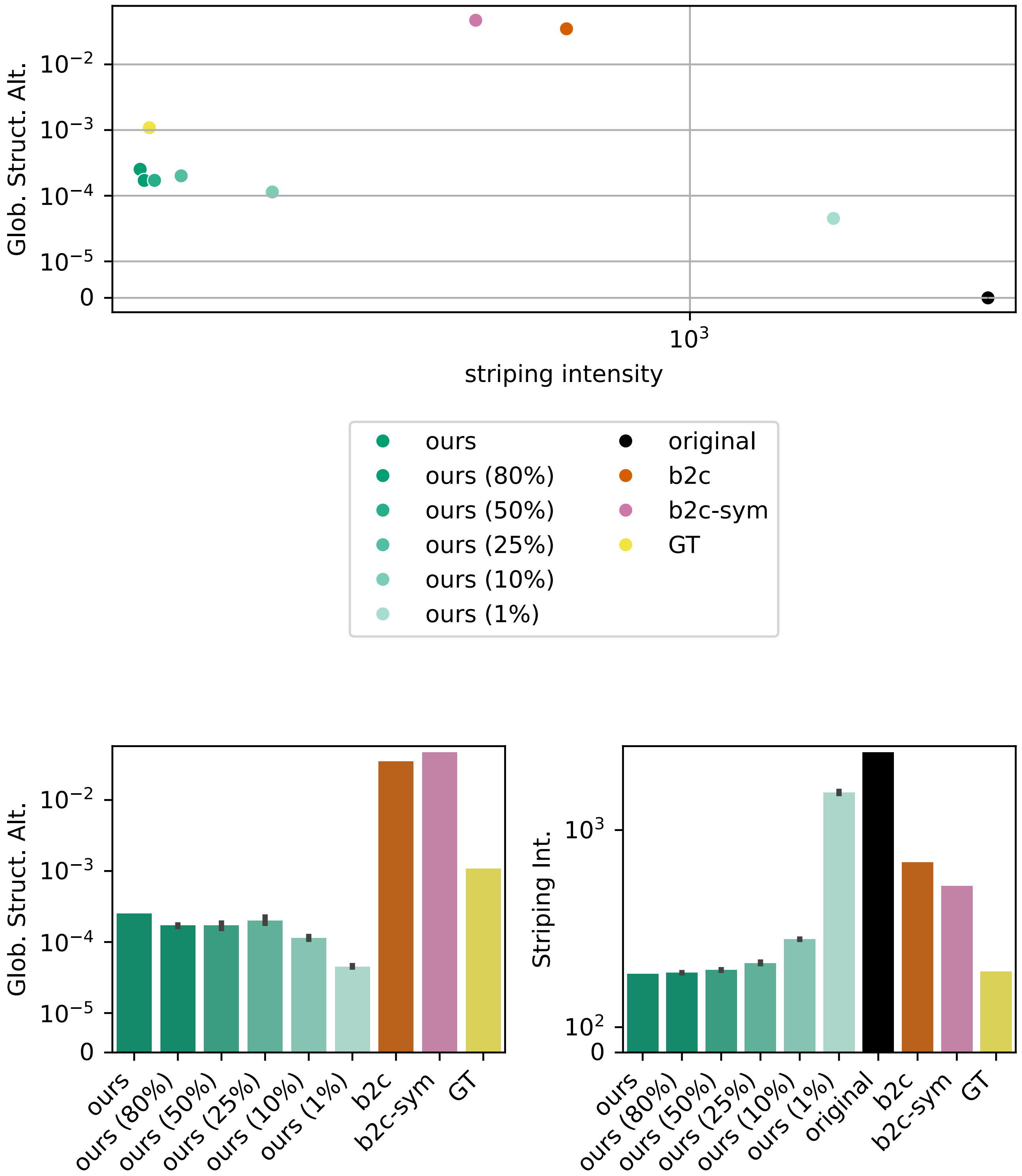


Fig. S36: **Missing cells experiment on the synthetic mouse brain dataset.** Global structure alteration score and striping intensity for different percentages of retained cells. Error bars indicate the standard deviation across three random seeds.

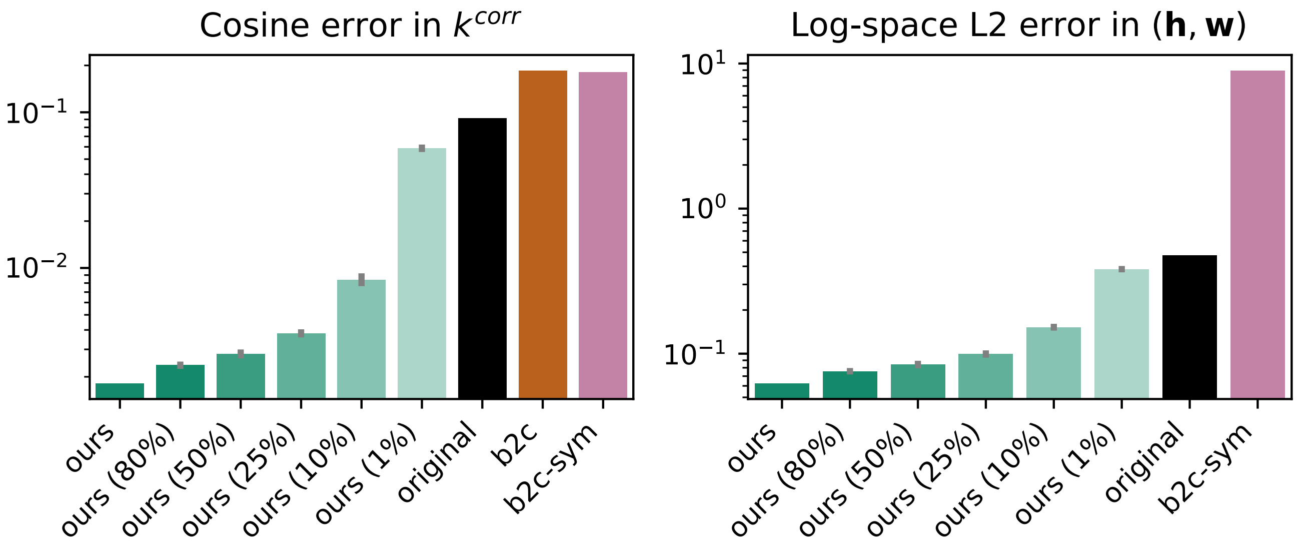


Fig. S37: **Missing cells experiment on the synthetic mouse brain dataset.** Cosine error in $k^{\mathrm{corr}}$ (left) and log-space $\ell_2$ error in $(\mathbf{h}, \mathbf{w})$ (right) for different percentages of retained cells. Error bars indicate the standard deviation across three random seeds.

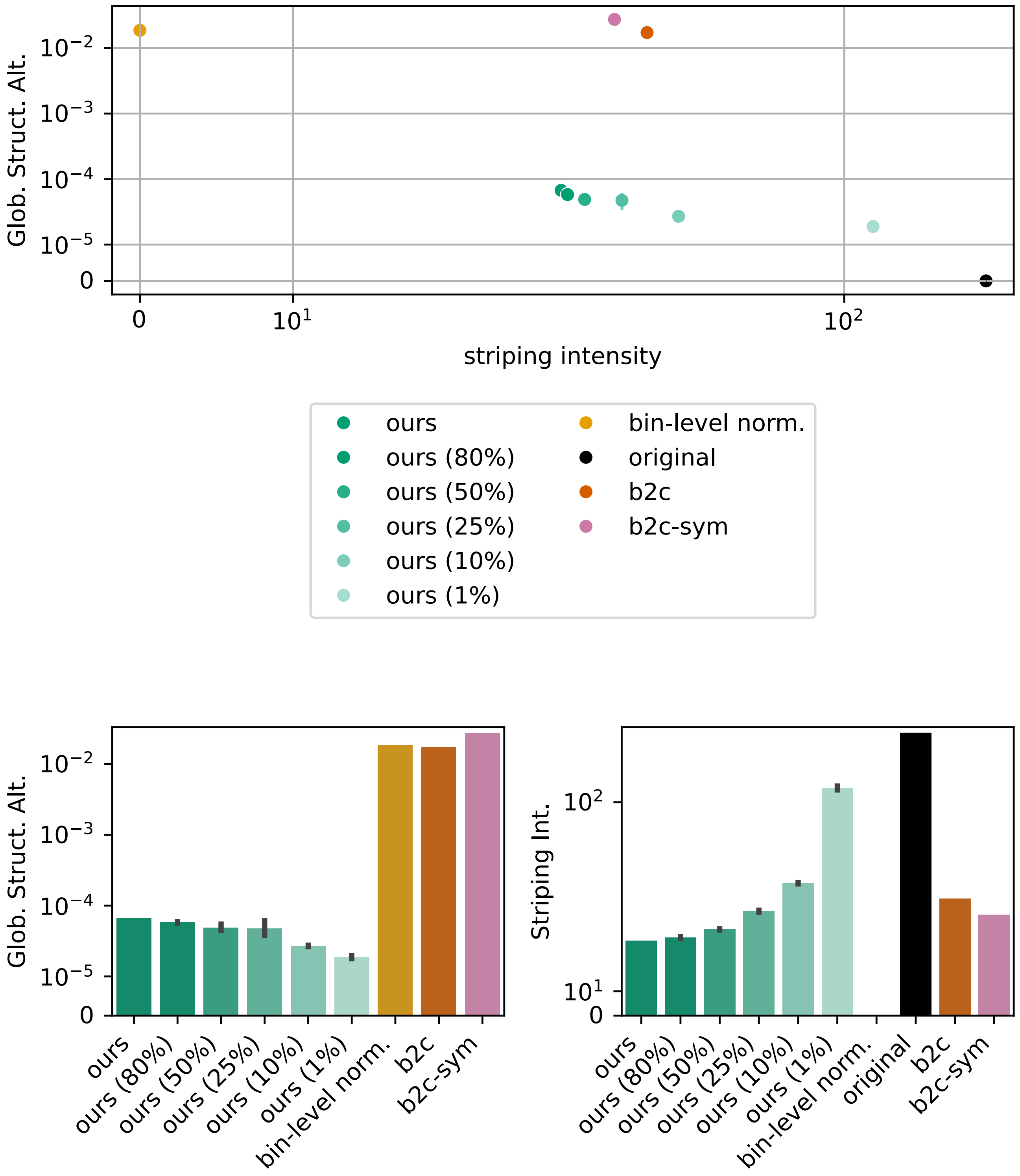


Fig. S38: **Missing cells experiment on the human lymph node dataset.** Global structure alteration score and cytoplasmic striping intensity for different percentages of retained cells. Error bars indicate the standard deviation across three random seeds.

#### S4.1.2. Oversegmentation

Our method is highly robust to oversegmentation in both the synthetic mouse brain and human lymph node datasets (Figures S39, S40 and S41). In both cases, performance remains satisfactory up to 90% of split cells. This robustness can be explained by the fact that, even at high levels of oversegmentation, the number of observed nuclei remains much larger than the number of model parameters to estimate. In this setting, the model is not fundamentally misspecified; rather, oversegmentation introduces additional parameters that are not strictly necessary. As a result, the method can still recover a satisfactory solution despite this increased model complexity.

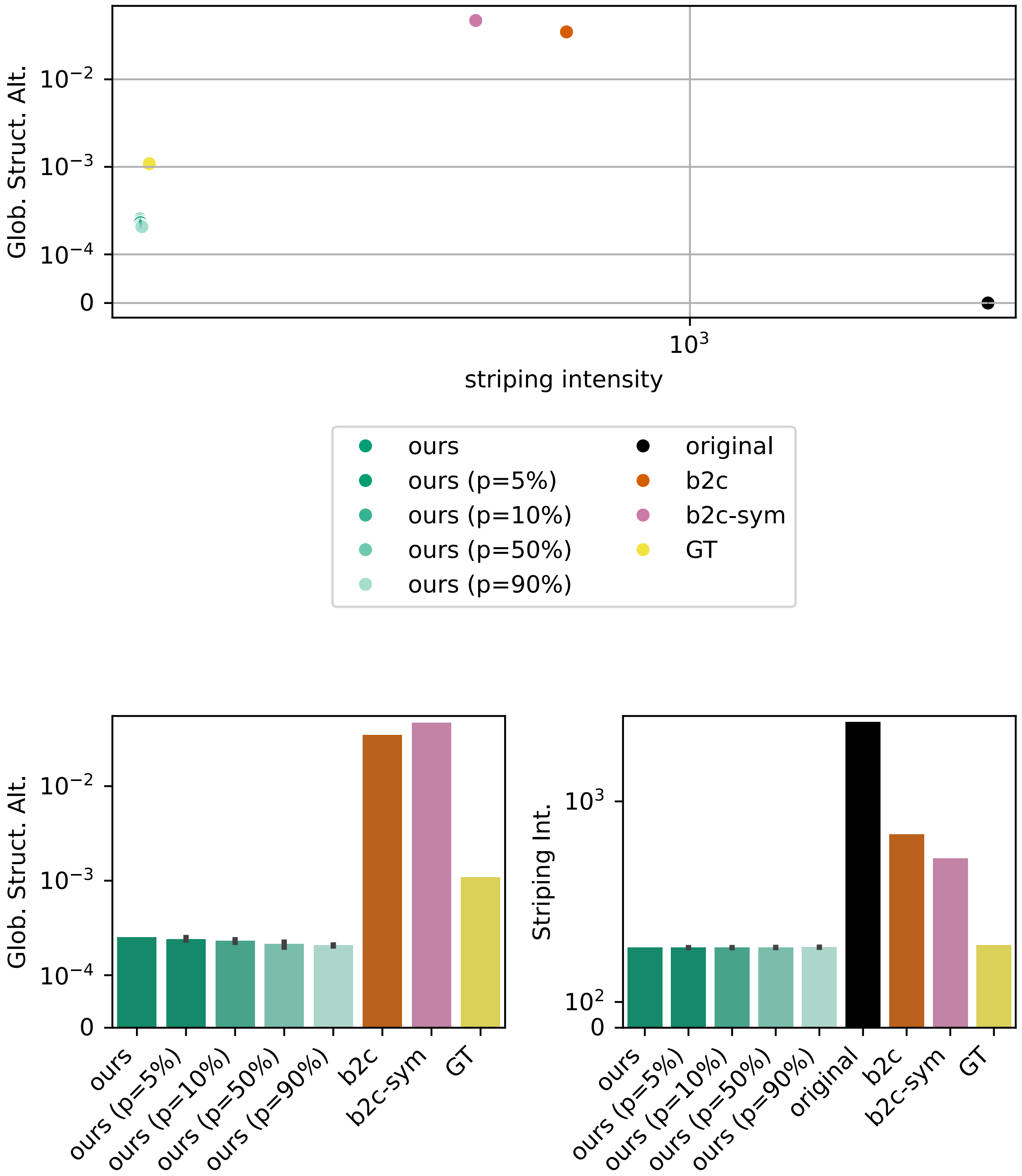


Fig. S39: **Oversegmentation experiment on the synthetic mouse brain dataset.** Global structure alteration score and striping intensity for different percentages of split cells. Error bars indicate the standard deviation across three random seeds.

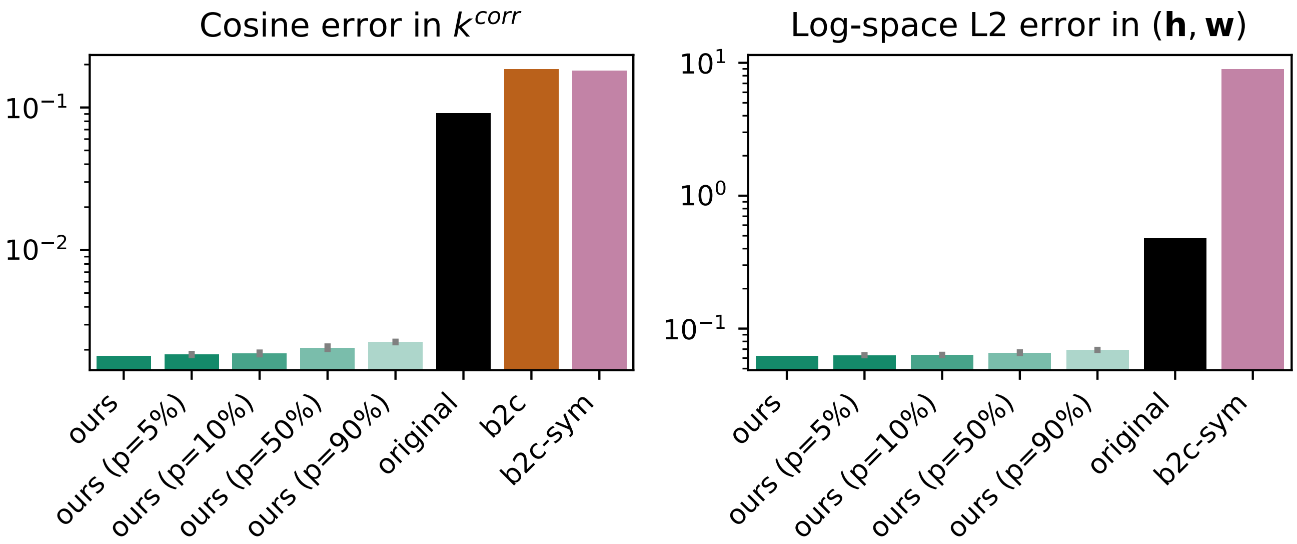


Fig. S40: **Oversegmentation experiment on the synthetic mouse brain dataset.** Cosine error in $k^{\text{corr}}$ (left) and log-space $\ell_2$ error in $(\mathbf{h}, \mathbf{w})$ (right) for different percentages of split cells. Error bars indicate the standard deviation across three random seeds.

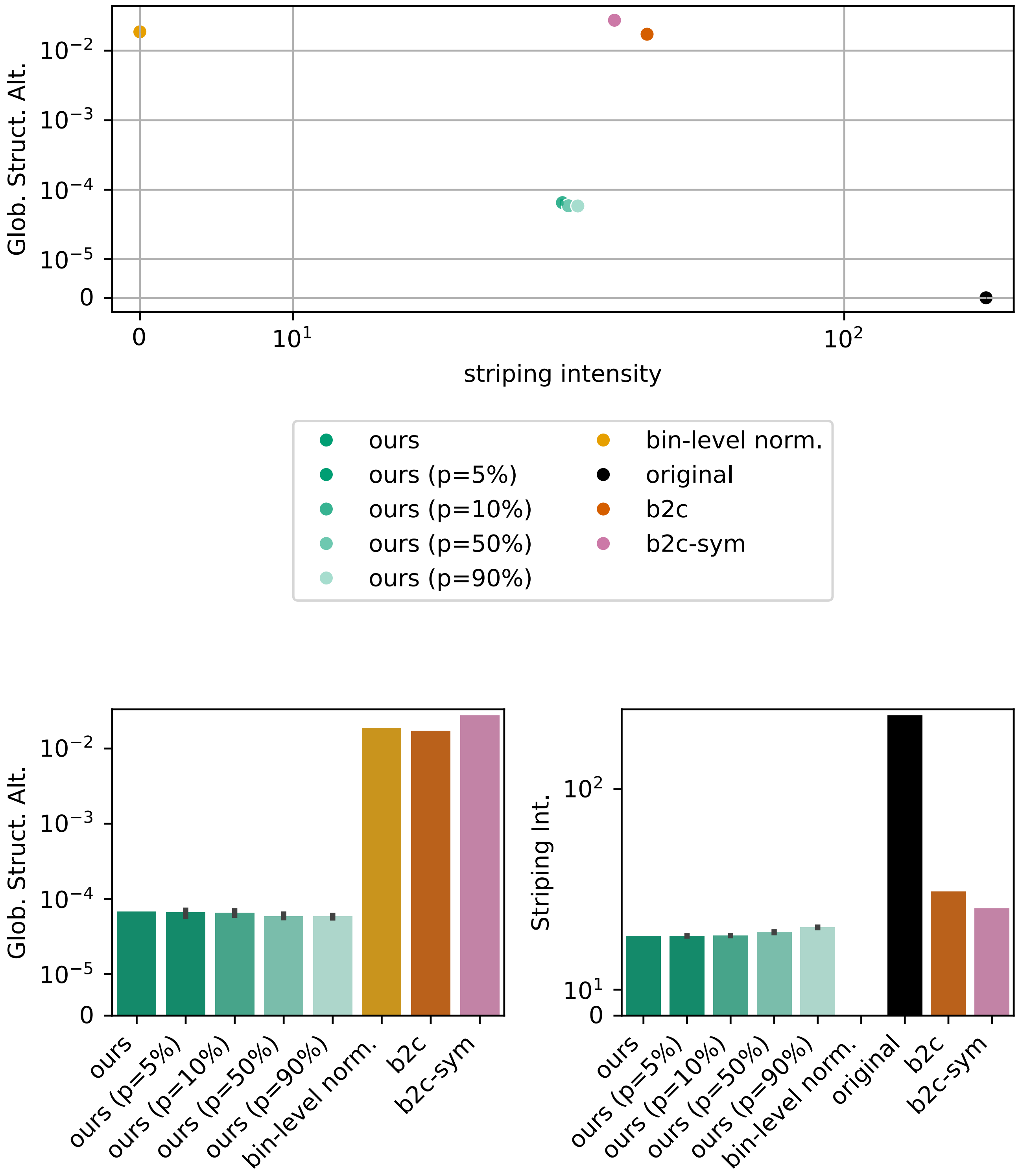


Fig. S41: **Oversegmentation experiment on the human lymph node dataset.** Global structure alteration score and cytoplasmic striping intensity for different percentages of split cells. Error bars indicate the standard deviation across three random seeds.

#### S4.1.3. Undersegmentation

Undersegmentation has a stronger impact on performance than missing cells or oversegmentation in both the synthetic mouse brain and human lymph node datasets (Figures S42, S43 and S44). On the synthetic data, our method remains competitive up to 25% of fused edges, outperforming the other baselines in terms of stripe factor estimation error and corrected count accuracy. Interestingly, at more extreme merging rates, striping intensity remains low while the global structure alteration score increases. The same trend is to be observed on the real human lymph node data for merging rates exceeding 50%. This suggests that the method is still able to remove the striping artifact, but at the cost of distorting the underlying count distribution. This behavior is not explained by the presence of isolated cells without edges (Figure S35), as the same trend is also observed for the **ours (collapsed label)** baseline. We also repeated the qualitative analysis shown in Figure 5 for the undersegmentation experiments. In agreement with the quantitative results, at high merging rates the striping artifact is visually removed in region b (Figure S45), whereas the count distribution is altered in region a (Figure S46).

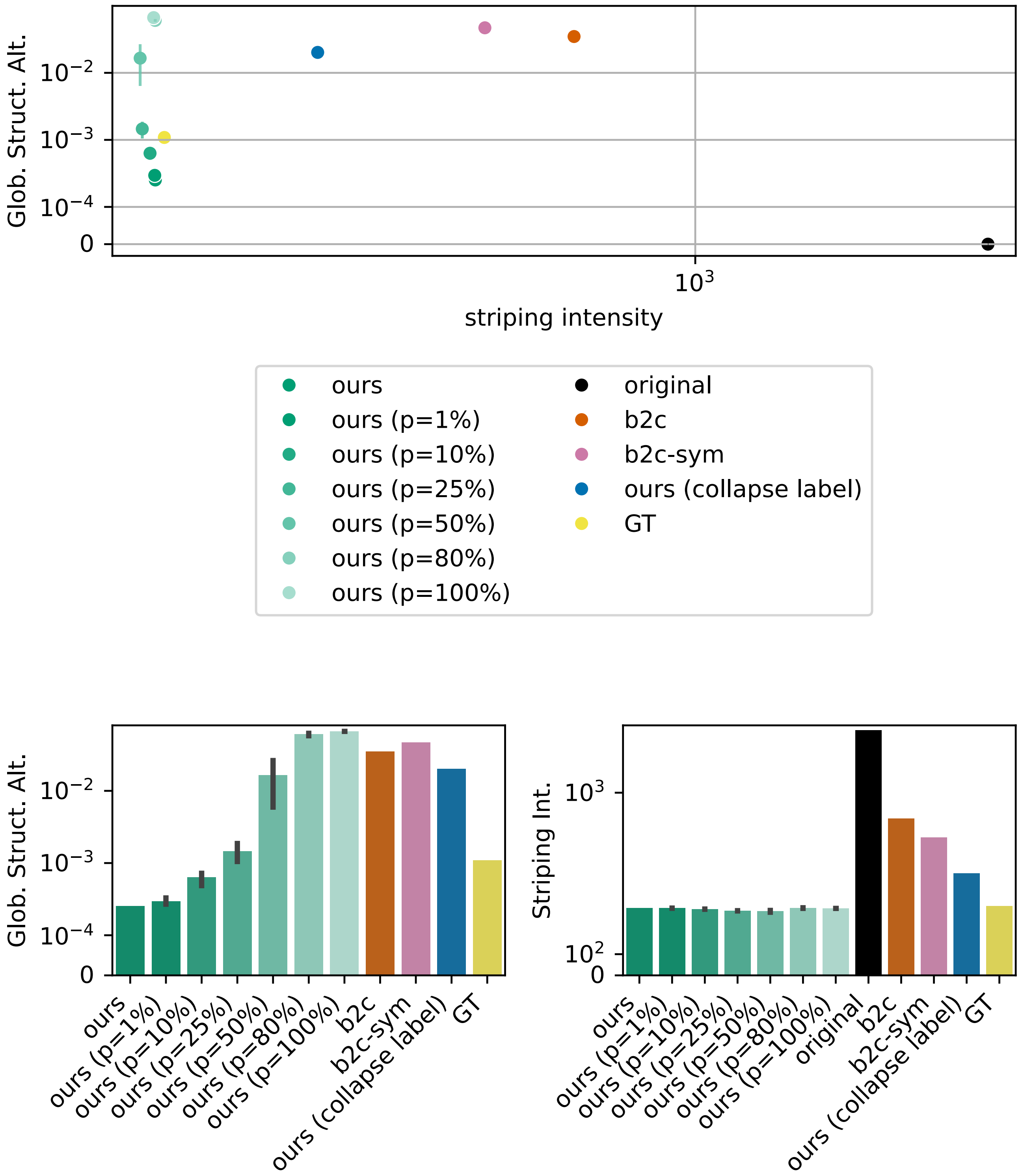


Fig. S42: **Undersegmentation experiment on the synthetic mouse brain dataset.** Global structure alteration score and striping intensity for different percentages of fused neighboring cells. Error bars indicate the standard deviation across three random seeds.

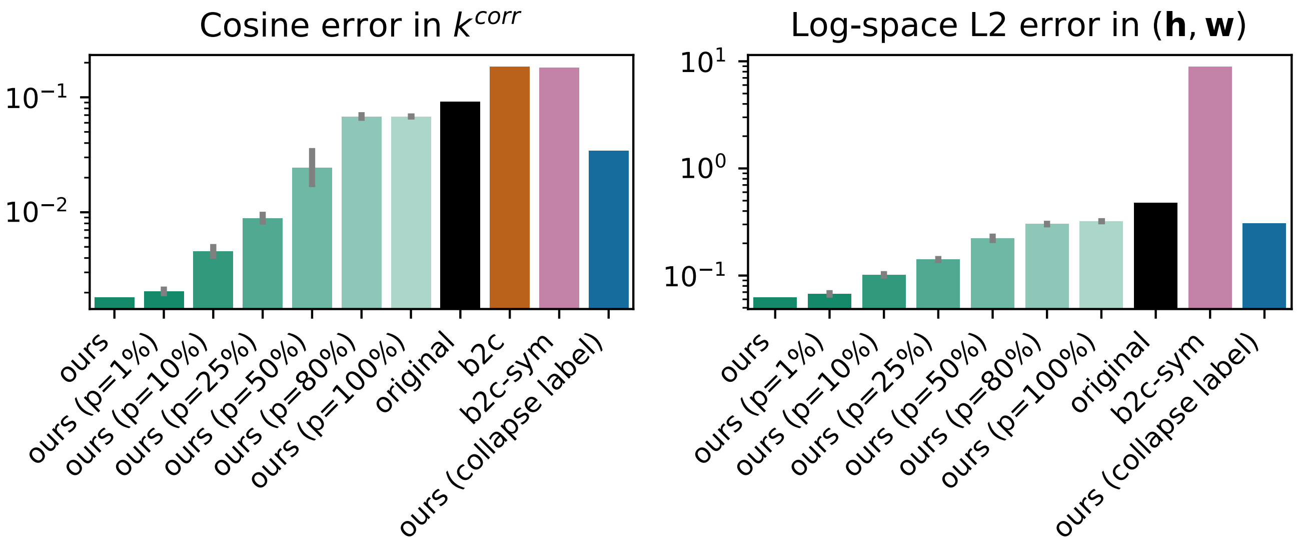


Fig. S43: **Undersegmentation experiment on the synthetic mouse brain dataset.** Cosine error in $k^{\text{corr}}$ (left) and log-space $\ell_2$ error in $(\mathbf{h}, \mathbf{w})$ (right) for different percentages of fused neighboring cells. Error bars indicate the standard deviation across three random seeds.

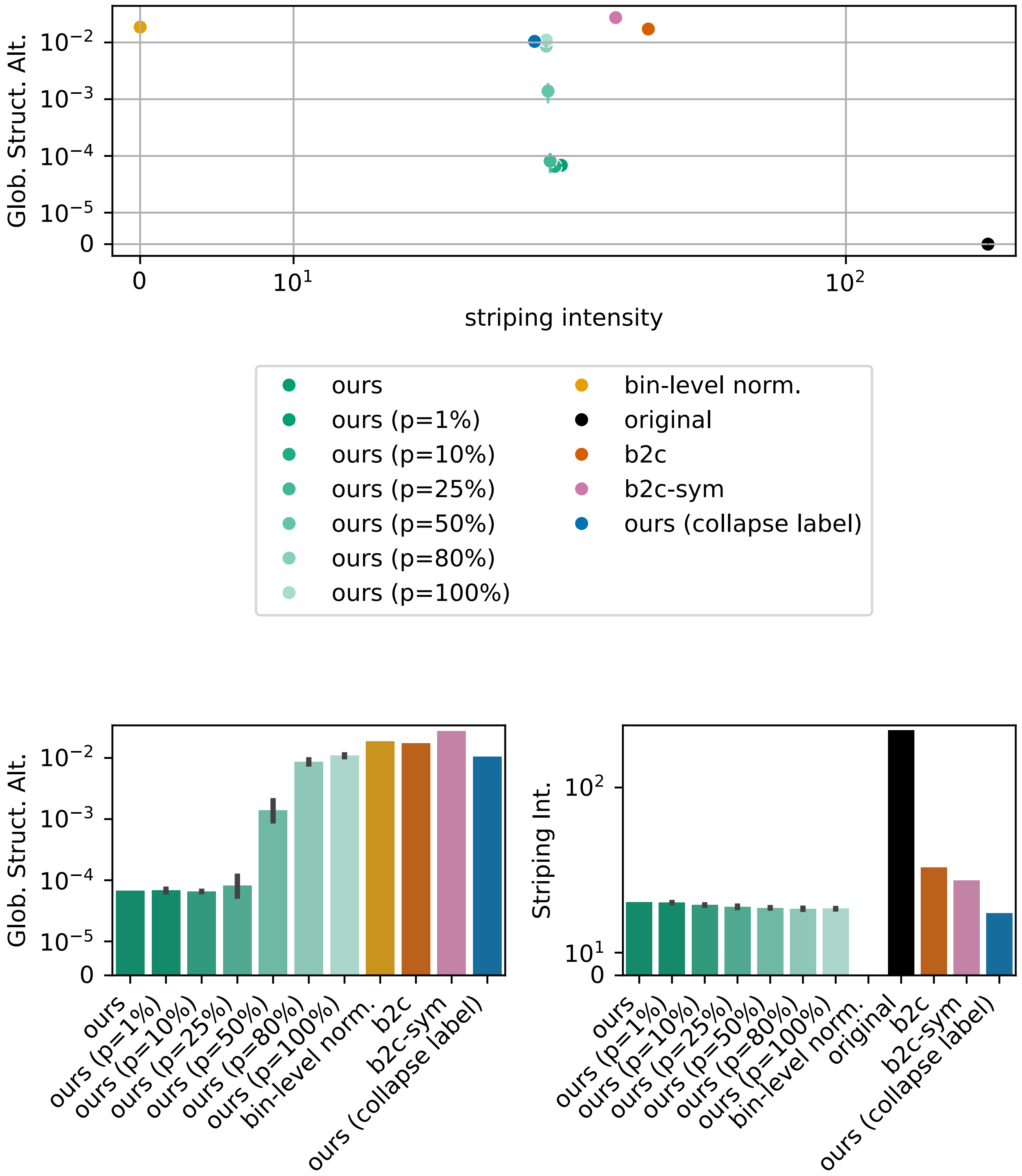


Fig. S44: **Undersegmentation experiment on the human lymph node dataset.** Global structure alteration score and cytoplasmic striping intensity for different percentages of fused neighboring cells. Error bars indicate the standard deviation across three random seeds.

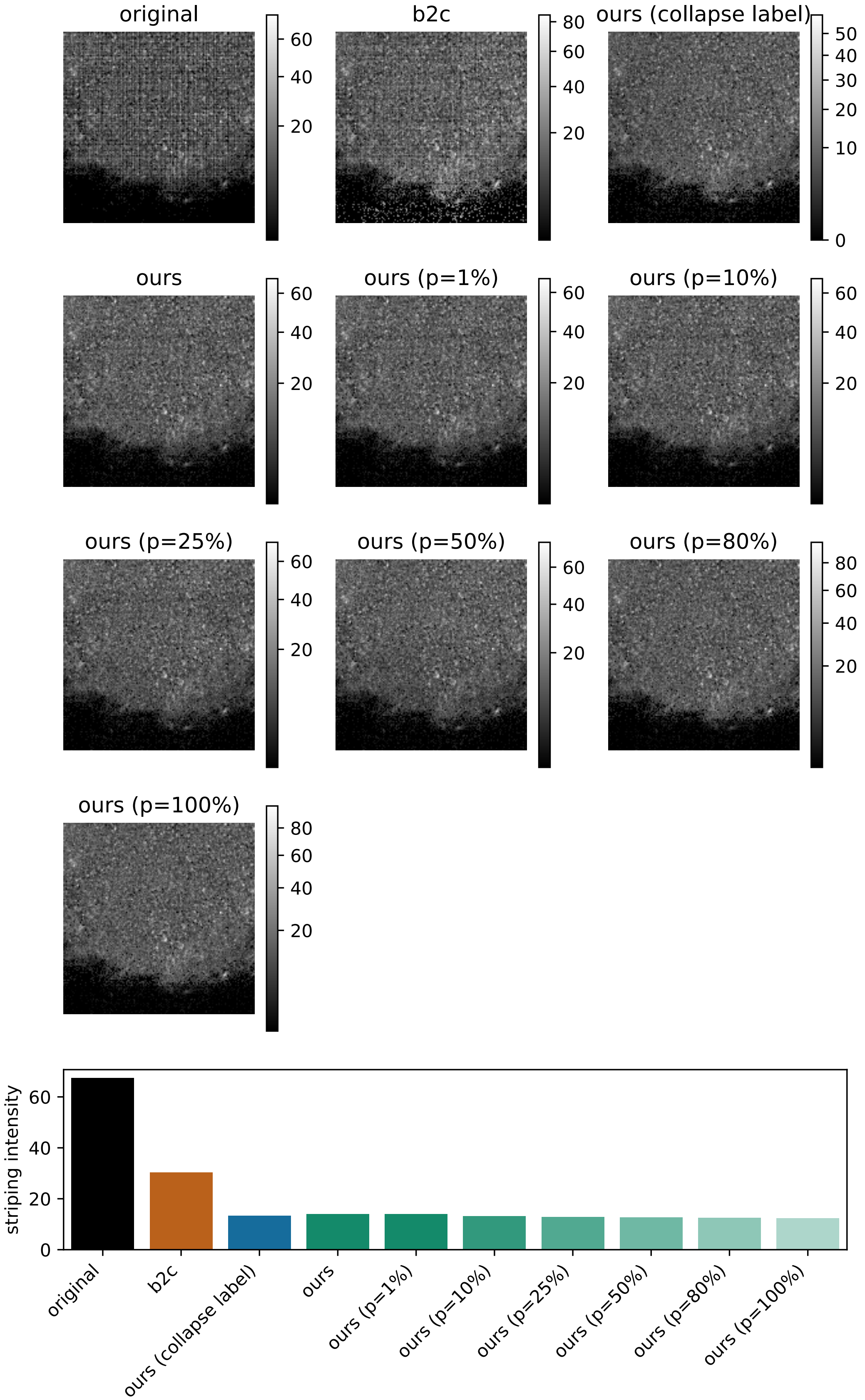


Fig. S45: **Undersegmentation experiment on the human lymph node dataset.** (top) Zoom on subregion b (see Figure 5) for different percentages of fused neighboring cells (for one seed). (bottom) Comparison of cytoplasmic striping intensity within subregion b.

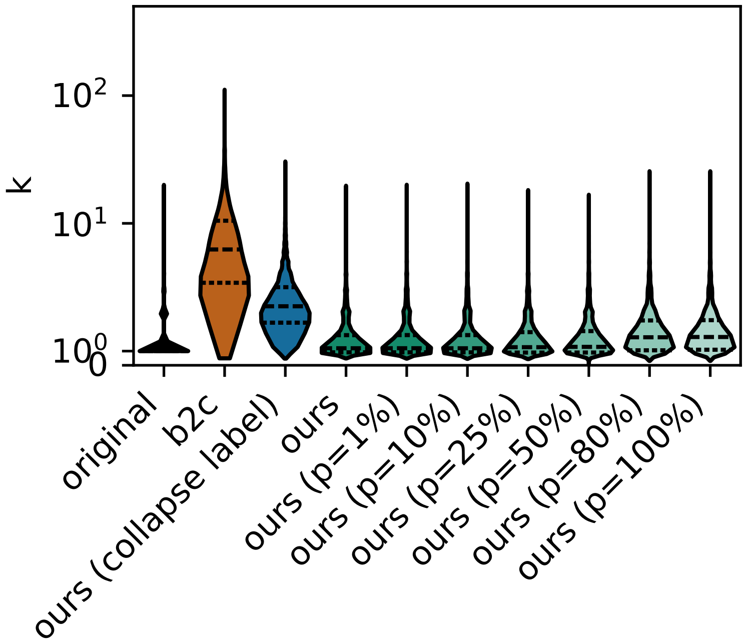


Fig. S46: **Undersegmentation experiment on the human lymph node dataset.** Count distribution in region a (see Figure 5) for different percentages of fused neighboring cells (for one seed).

#### S4.2. Summary

Overall, our method is extremely robust to missing-cell errors and oversegmentation, as the number of observed nuclei remains much larger than the number of parameters to estimate. It is also robust to moderate undersegmentation. At more extreme levels of undersegmentation, the method still removes the striping artifact, but tends to alter the global count distribution.

### S5. Evaluation of impact on downstream tasks

#### S5.1. Cell typing

We evaluated the impact of destriping on cell typing. To this end, we first aggregated bin-level gene expression to the nucleus level by summing counts across all bins assigned to the same nucleus. We then assigned cell types using `celltypist` on rescaled, log-transformed expression profiles, with the *Mouse Brain* reference model [Domínguez Conde et al., 2022].

Figure S48 shows that, on the real mouse brain dataset, cell-type assignments vary slightly depending on the destriping method, with pairwise agreement ranging from 84% to 91%. On synthetic data, we define ground-truth cell labels by applying `celltypist` to the expected gene expression under the generative model, i.e., before introducing the striping effect. Figure S47 shows that, on these simulated data, our method yields more accurate cell-type assignments than the other baselines. For reference, we also include the baseline GT-destriped, which corresponds to cell typing after destriping with the ground-truth stripe factors.

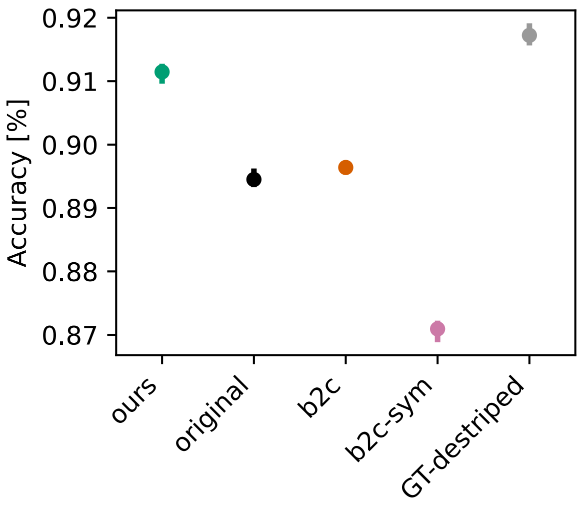


Fig. S47: **Cell typing results on the synthetic mouse brain dataset.** Accuracy of the inferred cell-type assignments. The reference labels are obtained by applying `celltypist` to the expected gene expression under the generative model, before introducing the striping effect. GT-destriped denotes the result obtained after destriping with the ground-truth stripe factors. Our method yields more accurate cell-type assignments than the baselines original, b2c, and b2c-sym, and approaches the performance of GT-destriped. Error bars indicate the standard deviation across the three random seeds used for data generation.

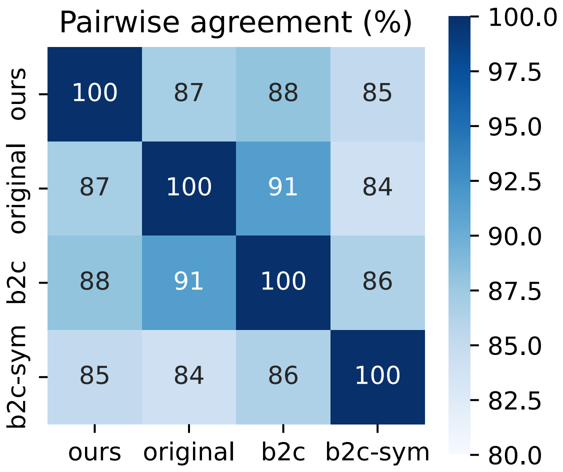


Fig. S48: **Cell typing results on the real mouse brain dataset.** Pairwise agreement between the cell-type assignments obtained after applying the different destriping methods.

### S5.2. Differential gene expression on the zebrafish head data

We evaluated the impact of destriping on downstream differential expression by comparing the eye and gill regions of the zebrafish head (Fig. S49). We first restricted the analysis to the 2,000 most highly variable genes, selected using the Seurat v3 method. Differential expression was then assessed with a Mann–Whitney U-test, and resulting $p$-values were corrected for multiple testing using the Benjamini–Hochberg procedure. Effect sizes are small in absolute value because VisiumHD data are highly sparse at the bin level.

At a false discovery rate of 5%, all methods yielded the same set of 1,273 significant genes. However, for approximately half of these genes, b2c and b2c-sym produced log fold-changes with opposite sign compared to the original data and our method (Fig. S50). Among these genes, *fgfr2* is of particular interest, as it has been reported to be highly expressed in the developing and adult zebrafish eye [Hochmann et al., 2012], consistent with the direction of the log fold-change obtained from the original data and our method. Figure S51 shows the nonzero expression of *fgfr2* in the two selected regions. In b2c and b2c-sym, artificially elevated values are introduced in the gill region, leading to an inversion of the estimated fold-change. This elevated expression in the gill is consistent with the artifactually high counts introduced by b2c and b2c-sym in the lower band of the slide, as observed in Fig. 4.

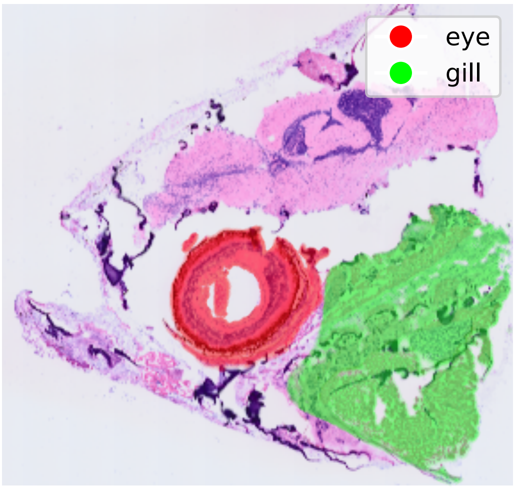


Fig. S49: **Regions used for differential expression analysis in the zebrafish head.** H&E staining image with the eye and gill regions highlighted.

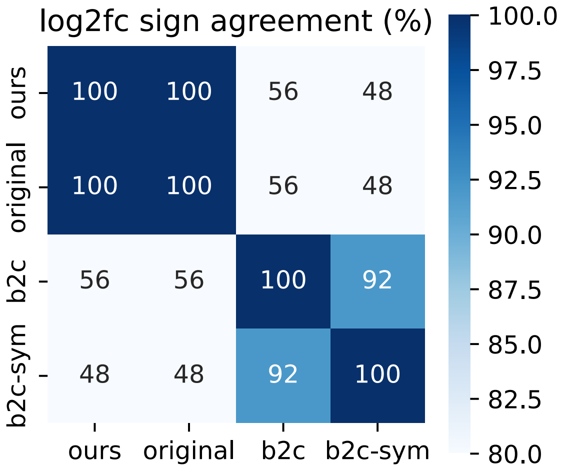


Fig. S50: **Agreement in differential-expression direction across destriping methods.** Pairwise agreement in the sign of the fold-change among significantly differentially expressed genes.

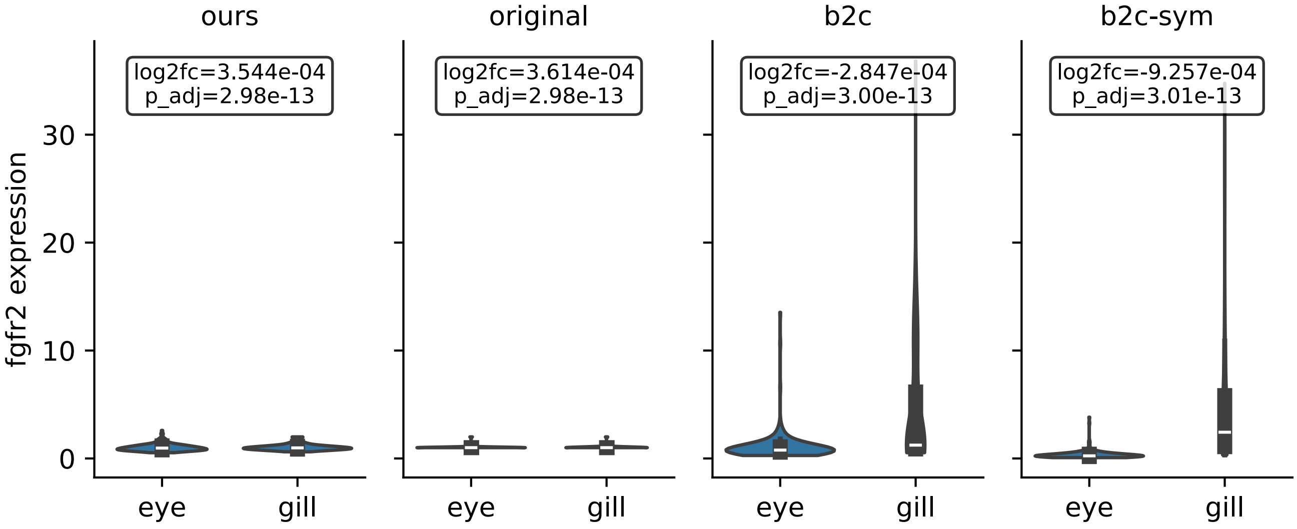


Fig. S51: **Expression of *fgfr2* in the eye and gill regions.** Distribution of nonzero *fgfr2* expression values for the different destriping methods. b2c and b2c-sym introduce artificially elevated values in the gill region. The top panel reports the $\log_2$ fold-change and corresponding $p$-value from the Mann–Whitney U-test. Consistent with the elevated gill expression, b2c and b2c-sym yield a reversed $\log_2$ fold-change relative to the original data and our method.

### S6. Computational requirements

| | # nuclei | ours (fitting) memory (GB) | ours (fitting) time (min) | ours (destriping) memory (GB) | ours (destriping) time (min) | b2c memory (GB) | b2c time (min) |
|---|---|---|---|---|---|---|---|
| mouse brain | 61842 | 1.26 | 8.44 | 5.19 | 0.44 | 2.61 | 0.07 |
| zebrafish head | 121779 | 0.72 | 8.73 | 1.18 | 0.12 | 1.19 | 0.02 |
| human lymph node | 292813 | 1.76 | 30.00 | 3.64 | 0.22 | 0.89 | 0.03 |
| mouse embryo | 448092 | 3.17 | 49.75 | 8.25 | 0.70 | 3.66 | 0.11 |

**Table S1.** Computational requirements of our method across four real datasets using four CPU cores.

When run on a modern Apple M1-based Mac, our method achieved an approximately 3× speedup relative to the runtimes reported in Table S1, and all datasets completed in less than 11 minutes.
